# Electrophysiological dissociation of human posterior cingulate cortex contributions to value- and memory-based decision-making

**DOI:** 10.64898/2026.08.12.744505

**Authors:** Seth R. Koslov, Hernan G. Rey, Sarah R. Heilbronner, Nicole R. Provenza, Sameer A. Sheth, Kathryn A. Davis, Han-Chiao I. Chen, Joseph W. Kable, Benjamin Y. Hayden, Brett L. Foster

## Abstract

The human posterior cingulate cortex (PCC) is routinely implicated in cognition and disease, yet its specific functional contributions remain unclear. Historically, human neuroimaging has linked the region to episodic memory and the default mode network, while a distinct non-human primate electrophysiology literature has focused on economic decision-making. Integrating anatomical evidence with these literatures, it has recently been proposed that this divergence reflects subregional organization, with dorsal PCC as a potential convergence site for value-based and memory-based decisions. Here, we recorded local field potentials (LFPs) and single units from human PCC while the same participants performed matched value- and memory-based decision tasks. LFPs in dorsal but not ventral PCC showed sustained engagement across both tasks, with risk sensitivity emerging only after the decision. In contrast, single-unit activity was more temporally circumscribed and could be grouped into response profiles active before or after the decision. Dorsal but not ventral PCC engagement further extended to memory encoding, recognition, and confidence judgments. Together, these findings reveal a consistent functional dissociation, identifying dorsal PCC as a domain-general interface between evaluative and mnemonic systems. In doing so, they align human and non-human primate accounts of PCC function and help orient future targeted studies of its role in cognition and disease.

## Introduction

The posterior cingulate cortex (PCC) has long attracted interest across basic and clinical human neuroscience, owing to its distinctly high resting metabolic activity (Gusnard & Raichle, 2001; Raichle et al., 2001), its position as a hub within large-scale brain networks (Gordon et al., 2018; Hagmann et al., 2008; Menon, 2023; Raichle, 2015), and its broad association with neurodegenerative and psychiatric diseases (Leech & Sharp, 2014; Maass et al., 2019; Vogel et al., 2021). Despite this prominence, the PCC’s specific functional contributions have remained difficult to elucidate, in part because the region has rarely been the primary target of investigation. Instead, its function has largely been characterized through neuroimaging studies of the default mode network (DMN) as a whole, leaving the PCC’s unique role difficult to isolate. However, contemporary neuroimaging work has shown that the PCC participates in multiple large-scale networks, and correspondingly, a diverse set of cognitive functions extending beyond those typically ascribed to the DMN (Bzdok et al., 2015; DiNicola et al., 2023; Du et al., 2024; Gilmore et al., 2015; Rolls et al., 2023; Vincent et al., 2008). This heterogeneity has recently been proposed as a way to address a longstanding divergence in the literature, in which human neuroimaging research emphasized the PCC’s role in episodic memory, while non-human primate (NHP) electrophysiology has emphasized its role in decision-making (Foster et al., 2023). Given the lack of a rodent PCC homolog (Vogt & Paxinos, 2014), reconciling this functional distinction across primate species and recording techniques is an important and compelling avenue for progress in understanding PCC’s role in cognition and disease.

The apparent tension between human neuroimaging and NHP electrophysiological accounts of PCC function may be reconciled by considering prior evidence of the distinct functional roles of the dorsal and ventral subregions of PCC. Moving beyond a unitary functional view of the entire PCC, precision connectivity studies in humans have associated dorsal PCC (dPCC) with frontoparietal control and salience network regions, while linking ventral PCC (vPCC) to DMN regions (Gordon, Laumann, Gilmore, et al., 2017; Kwon et al., 2025; Vincent et al., 2008; Willbrand et al., 2022). Strikingly, a complementary dissociation also exists within human neuroimaging work examining differing episodic remembering tasks. Item-recognition decisions are uniquely associated with dPCC activity, while autobiographical retrieval is predominantly associated with vPCC activity (Chen et al., 2017; McDermott et al., 2009). Furthermore, human neuroimaging work has frequently implicated dPCC in decision-making and cognitive control processes (Aponik-Gremillion et al., 2022; Corlett et al., 2022; Jauhar et al., 2021; Kable & Glimcher, 2007; Levy & Glimcher, 2011), though these observations have received considerably less attention and subsequent functional specification than parallel work in regions like the anterior cingulate cortex. In light of this human literature, it is important to note that electrophysiological recordings in NHP PCC have primarily targeted an area homologous to human dPCC, with a resulting focus on the region’s role in decision-making (Barack et al., 2017; Hayden et al., 2008; McCoy & Platt, 2005). Together, these lines of evidence support an important dorsal-ventral subdivision within PCC, with converging findings highlighting the engagement of dPCC during decision-making and suggesting that the subregion’s episodic memory associations reflect decision-oriented behaviors (Foster et al., 2023). However, this proposed role for dPCC at the intersection of decision-making and memory processes has been inferred across separate research domains, species, and recording methods, and has not yet been tested directly.

In seeking to better align the human and NHP PCC literatures, it is constructive to consider the open questions within each subfield, and how they relate to obstacles inherent to their respective approaches. In human studies, predominantly using functional magnetic resonance imaging (fMRI), outstanding questions relate to limitations in the temporal resolution of signals, resulting in ambiguity about when PCC engagement occurs relative to decision behavior. For example, human fMRI studies differ on whether they link dPCC activity to pre-decisional deliberation or to post-decision monitoring processes (Bartra et al., 2013; Clithero & Rangel, 2014; Oldham et al., 2018). As noted above, while there have been early associations of PCC with value computations (Engelmann & Tamir, 2009; Kable & Glimcher, 2007), a prominent literature has associated the region with recognition decisions (Elman et al., 2013; Gilmore et al., 2019; Kim, 2013; Koslov et al., 2024), and more recently, with memory-guided decisions that require episodic retrieval or temporally extended information integration (Kolling et al., 2014; Rosen et al., 2016; Visalli et al., 2019). Although both value- and memory-based decisions can be readily studied in humans, they have rarely been examined together, developing as largely separate literatures that leave open whether dPCC engagement generalizes across decision types. In contrast, studies of NHP PCC predominantly use electrophysiology, focusing on single-unit responses, yielding temporally specific predictions of PCC responses during post-decision monitoring and behavioral policy updating (Barack & Platt, 2021; Heilbronner et al., 2011; Heilbronner & Platt, 2013; Pearson et al., 2009). However, unlike human studies, where directly comparing PCC responses during different types of value- and memory-based decisions is feasible, implementing complex multi-task paradigms involving episodic memory in non-human species remains challenging. Resolving these open questions of where and when PCC is engaged during a decision, and how this generalizes across decision types, requires within-subject, multi-scale electrophysiological recordings during both value- and memory-based decisions, a combination uniquely achievable through human intracranial recordings.

To address these core questions about PCC function, we leveraged invasive intracranial recordings to measure local field potentials (LFPs) and single-unit activity from human participants while they performed matched value- and memory-based decision-making tasks. Consistent with the subregional framework, broadband gamma (BBG; 70 - 150 Hz) LFP responses revealed robust population-level engagement of dPCC, but not vPCC. For both value- and memory-based decisions, dPCC BBG responses demonstrated a similarly sustained response profile, spanning trial onset through feedback presentation. Interestingly, risk- and value-related modulation of BBG responses emerged selectively during the post-decision period. Unlike the sustained population-level BBG response, single-unit activity was more temporally circumscribed, with dissociable clusters of units engaged during specific decision-making stages. Furthermore, consistent response profiles across tasks were observed only for a subset of units, while others demonstrated task-specific engagement. These observations underscore the value of multiscale recordings for understanding heterogeneous cortical regions like PCC, with population-level and single-unit measures providing complementary information about contributions to decision-making. Hippocampal recordings, collected for comparison given the region’s established role in memory and decision-making (Bakkour et al., 2019; Biderman et al., 2020; Lopez-Persem et al., 2020), revealed response profiles broadly mirroring those of dPCC. Together, these results provide direct electrophysiological evidence identifying dPCC as a convergence site for evaluative and mnemonic decisional processes, resolving key ambiguities about the timing and generalizability of its involvement in decision-making. In doing so, this work reconciles disparities between human and NHP accounts of PCC function and supports a framework for better understanding this enigmatic region’s broader role in cognition and disease.

## Results

### Intracranial PCC recordings during decision-making

Electrophysiological activity in human PCC was recorded using stereo-electroencephalography (sEEG) depth probes in 20 participants undergoing invasive monitoring for epilepsy (female = 12, male = 8). No included participants had a clinically determined epileptogenic focus within the PCC. Across participants, 48 PCC recording sites were included in analyses, with 25 located in the dorsal PCC (dPCC) and 23 located in the ventral PCC (vPCC) (Figure 1a). A subset of probes contained a bundle of 8 microwires extending from the distal tip (yellow outlines, Figure 1a), allowing for the recording of single-unit spiking activity (participants n = 10; microwire sites n = 10, units n = 110). To characterize PCC involvement in decision-making (DM) processes, population-level local field potentials (LFPs) and single-unit spiking activity were recorded while participants performed a value-based DM task and a three-part memory-based DM paradigm (Figure 1b, see methods). Participants first performed the value-based DM task, where they attempted to earn points by selecting between a ‘safe’, guaranteed low-value reward or a ‘risky’ probabilistic chance at a higher-value reward. Next, participants completed a memory-based DM task, which was equated in structure and timing to the value-based DM task. DM tasks differed only in the source of information available for influencing decisions, expected value for the value-based task and memory encoding strength for the memory-based task, allowing neural responses to be characterized within a common framework (Figure 1c, see methods for details). To control and test memory strength, the memory-based DM task was preceded by an incidental encoding 1-back task (Figure 1e) and followed by a delayed-recognition task (Figure 1f).

**Figure 1.**
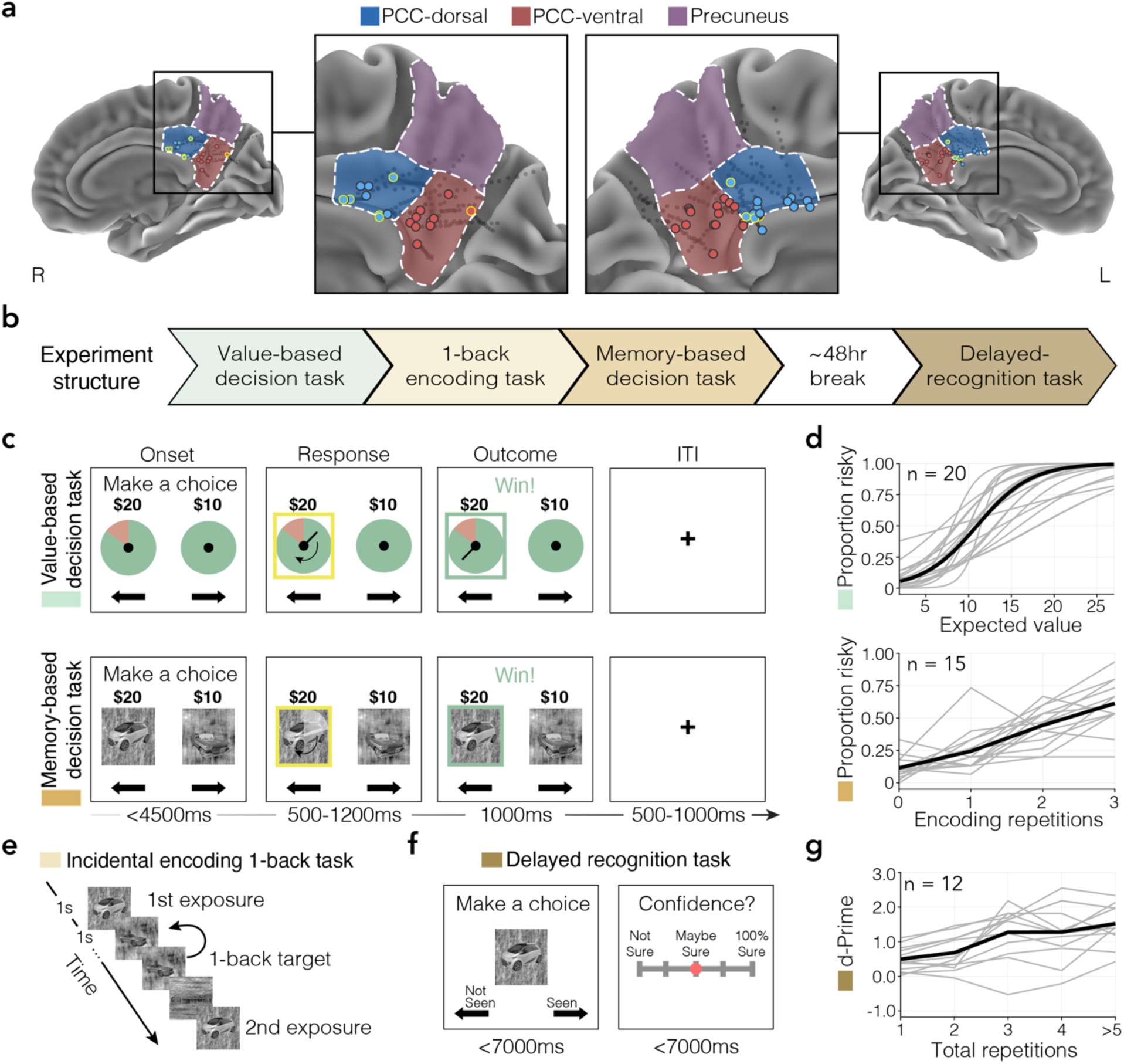
Anatomy, experimental design and task behavior. **a)** Normalized anatomical locations of intracranial recording sites across participants shown on the medial cortical surface of the MNI152 template. Insets and colors highlight medial parietal cortex (MPC) anatomy and corresponding electrode location designations (blue = dorsal PCC, red = ventral PCC, purple = precuneus). Medial electrodes from probes containing microwires are indicated with yellow outlines. **b)** Experimental task structure involved participants performing both a value-based decision-making task and a three-part memory paradigm occurring over multiple sessions/days during invasive recordings. First, participants performed a value-based decision-making task. Next, participants performed an incidental encoding 1-back task, followed by a memory-based decision-making task, and after a delay (∼48 hrs), a recognition memory task. **c)** The value-based (top) and memory-based (bottom) decision-making tasks had matched trial designs. For both tasks, participants attempted to earn points by selecting between risky and safe options. Across tasks, the safe option value was always $10 while the risky option was a higher amount that varied across trials. For the value-based task, reward probability was explicitly indicated by the proportion of green on the wheels (safe: 100%, risky: 20-90% in 10% increments). In the memory-based task, reward probability depended on memory for stimuli, with rewards delivered for selecting “old” stimuli and no reward for selecting “new” stimuli. The safe option was always a stimulus that had been frequently repeated during encoding, while the risky option could either be a novel (“new”) or infrequently repeated (“old”) stimulus. For both tasks, following trial onset, participants made a choice (<4500ms), followed by a post-response delay period (500-1200ms), veridical feedback (1000ms), and an inter-trial interval (ITI, 500-1000ms). **d)** Behavioral performance for individuals (grey lines) and across all participants (black lines) on both decision-making tasks is shown, confirming that the likelihood of selecting the risky option (y-axis) varied as a function of its expected value (top; value-based) or the number of incidental-encoding repetitions (bottom; memory-based). **e)** Participants completed an incidental 1-back encoding task in which stimuli were presented (1000ms each) either frequently (5 times) or infrequently (1-3 times) over spaced intervals, parametrically varying encoding strength for stimuli used in the memory-based decision-making task. Participants were instructed to press a button upon detecting a repeated image (1-back), but were not informed that stimuli would be used in the following memory-based decision-making task. **f)** For the delayed recognition task, participants were asked to make a seen/unseen (“old”/“new”) decision about individually presented stimuli (<7000ms, left). Recognition decisions were followed by a 5-point scale prompt where participants indicated decision confidence (<7000ms, right). **g)** Behavior on the delayed-recognition task for individuals (grey lines) and across all participants (black line) shows that memory performance (d-prime, y-axis) was modulated by the number of total stimulus repetitions across the entire experiment.

We first assessed whether participant behavior was impacted by the experimentally manipulated decisional factors, expected value and memory strength. For the value-based DM task, the proportion of trials participants selected the risky option increased as the expected value (reward value * reward probability) increased (z = 8.893, β = 0.317, 95% CI = [0.247, 0.387], p < 0.001, Figure 1d-top), behavior consistent with previous reports of risky decision-making in healthy controls and patient populations (Saez et al., 2018). In the memory-based DM task, the proportion of trials participants selected the risky option increased as a function of the number of times that they had seen the stimulus during the prior encoding task (t_(44)_ = 10.248, β = 0.169, 95% CI = [0.137, 0.201], p < 0.001, Figure 1d-bottom). The delayed-recognition task provided a traditional memory assessment to confirm stimulus encoding and evaluate long-term retention of stimuli. After a delay period (mean = 47.4 hrs), participants successfully recalled previously presented stimuli (t_(11)_ = 6.769, d’ = 0.830, 95% CI = [0.560, 1.100], p < 0.001) and were more accurate at remembering stimuli as a function of total repetitions (t_(47)_ = 7.204, β = 0.264, 95% CI = [0.193, 0.336], p < 0.001; Figure 1g). Together, these results confirm that decision-making behavior was systematically influenced by the experimentally manipulated decisional factors, expected value and memory encoding strength, validating the task design for subsequent neural analyses.

### Subregional PCC LFP responses during decision-making

To examine PCC contributions to decisional processes, we analyzed LFPs from macro electrodes in the dorsal and ventral PCC subregions (Figure 2a). We focused on broadband gamma (BBG; 70-150Hz) activity as it provides a time-resolved measure of coincident local population spiking dynamics (Foster et al., 2016; Miller et al., 2014; Ray & Maunsell, 2011). Analyses were performed using linear mixed-effects regression, including a random intercept term for participant to control for multiple recording sites within individuals (see methods for details). For concise reporting of statistical results, p-values are reported here with full regression statistics provided in supplementary tables. To characterize responses identified in a priori-determined time windows with respect to the broader temporal response profile, and to control for multiple comparisons across time, we also performed complementary FWER-corrected time-series analyses. In cases where a priori effects did not survive cluster-correction, FWER-corrected p-values are also provided in text. For both the value- and memory-based DM tasks, dPCC BBG activity was significantly elevated following trial onset and through the response period, returning to baseline following feedback (value-based, onset: p < 0.001; response: p < 0.001; feedback: p < 0.001; memory-based, onset: p = 0.004; response: p = 0.008; feedback: p = 0.004; Figure 2b, Table S2). Conversely, vPCC BBG activity was only significantly above baseline briefly during the value-based DM task trial onset period, but remained at baseline levels for all other time periods across both tasks (value-based, onset: p = 0.037; response: p = 0.451; feedback: p = 0.064; memory-based, onset: p = 0.168; response: p = 0.473; feedback: p = 0.145; Figure 2b, Table S2). Recruitment of dPCC, but not vPCC, during DM is consistent with observations of dPCC involvement in executive processing (Aponik-Gremillion et al., 2022; Kolling et al., 2014; Leech et al., 2011; Waskom et al., 2017). However, the functional organization of PCC has been shown to be highly individualized (Du et al., 2024; Gordon, Laumann, Adeyemo, et al., 2017; Koslov et al., 2024), such that anatomical labeling of recording sites may obscure meaningful functional distinctions, motivating a complementary data-driven clustering approach to characterizing PCC contributions to decision-making.

**Figure 2.**
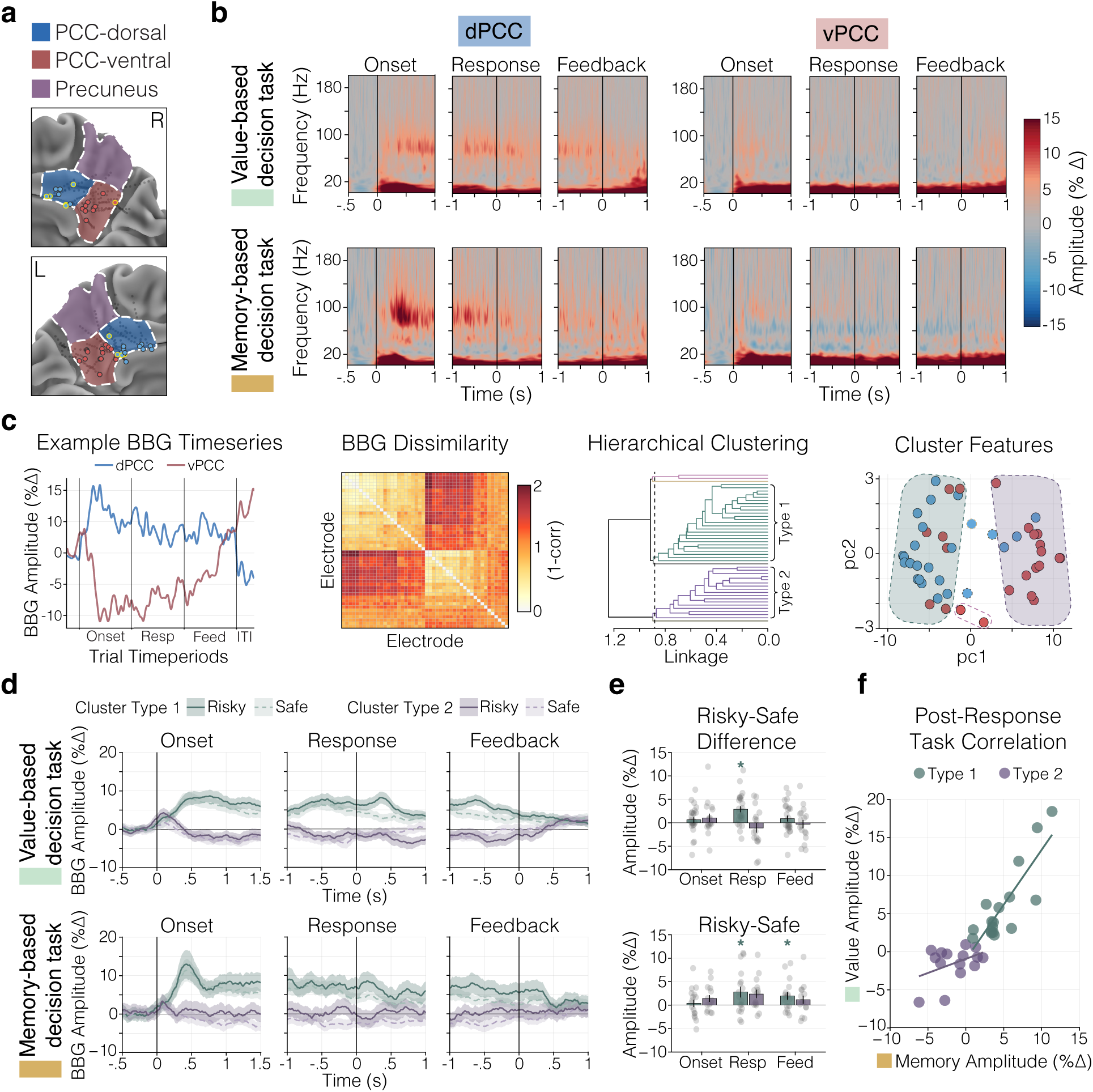
PCC LFP responses and clustering of response types during decision-making. **a)** Focused view of MPC, highlighting anatomy and recording site locations (macro electrodes n = 48, blue = dorsal PCC, red = ventral PCC, purple = precuneus). Medial electrodes from probes containing microwires are indicated with yellow outlines. **b)** Group-mean time-frequency spectrograms of local field potential (LFP) amplitude (percent signal change from pre-trial baseline; %Δ) dynamics during the value-based (top) and memory-based (bottom) decision-making tasks from recording sites located in the dorsal (blue, left) and ventral (red, right) PCC. Responses are aligned to trial onset (left), decision button-press response (center), and feedback presentation (right). **c)** Data-driven clustering approach for analysis of LFPs (left to right): i) baseline-corrected broadband gamma (BBG, 70-150 Hz) amplitude was concatenated across trial epochs into a single timeseries for each PCC electrode (blue = dPCC, red = vPCC); ii) a pairwise dissimilarity matrix (1-r) was computed between each electrode’s mean concatenated timeseries; iii) hierarchical clustering of the dissimilarity matrix yielded two primary clusters: type 1 (green) and type 2 (purple); iv) projection of dorsal (blue) and ventral (red) electrodes onto the first two principal components of the dissimilarity matrix, with colored outlines delineating the separation of cluster types in principal component space (green = type 1, purple = type 2). **d)** Group-mean BBG (70-150 Hz, percent signal change from pre-trial baseline) on risky (solid lines) and safe (dashed lines) trials for the value-based (top) and memory-based (bottom) tasks from electrode clusters 1 (green) and 2 (purple). Responses are aligned to trial onset (left), decision button-press response (middle), and feedback presentation (right). **e)** The mean difference in BBG response amplitude for risky minus safe trials during the 500 ms period following trial onset, response, and feedback time periods for electrode clusters 1 (green) and 2 (purple). **f)** Across-task correlation of mean-BBG amplitude during the post-response period (500 ms) between the value-based (y-axis) and memory-based (x-axis) tasks for type 1 (green) and type 2 (purple) electrodes. BBG amplitude was positively correlated across tasks for type 1 but not type 2 electrodes.

#### Clustering reveals distinct PCC LFP response types

For data-driven clustering (see methods for details), a mean BBG time series was computed for each site by concatenating activity across trial epochs from one value-based DM task block (Figure 2c, left), and then pairwise dissimilarity was calculated across all sites (Figure 2c, center-left). Hierarchical clustering of the resulting dissimilarity matrix yielded five unique clusters (Figure 2c, center-right; see Methods), with most sites belonging to cluster type 1 (green) or type 2 (purple). Projection of recording sites onto the first two principal components of the dissimilarity matrix (Figure 2c, right) revealed a clear separation between type 1 (green) and type 2 (purple) sites, with dPCC (blue) sites predominantly belonging to type 1 (76%, 19/25) and vPCC sites to type 2 (61%, 14/23, χ^2^(2, n = 48) = 12.795, p = 0.002, V = 0.516). Although the majority of dorsal and ventral PCC sites were segregated into distinct cluster types, some overlap was observed (3 dPCC sites in type 2; 7 vPCC sites in type 1; 3 dPCC sites and 2 vPCC sites designated as types 3-5), underscoring the limitations of purely anatomical classification and the utility of a data-driven clustering approach.

#### Dissociable involvement and risk sensitivity across PCC LFP response types

Having established two primary functional PCC cluster types, we next sought to characterize how each type was recruited during value- and memory-based decision-making. To do so, we first evaluated BBG responses from type 1 and type 2 sites across trial time periods (onset, response, and feedback). Type 1 sites demonstrated significantly elevated BBG activity from trial onset through the decision button-press response, returning to baseline after the feedback presentation period (value-based, onset: p < 0.001; response: p < 0.001; feedback: p < 0.001; memory-based, onset: p = 0.001; response: p < 0.001; feedback: p < 0.001; Figure 2d, green, Table S3a). We did not observe any differences from baseline for type 2 activity across task periods (value-based, onset: p = 0.819; response: p = 0.124; feedback: p = 0.569; memory-based onset: p = 0.642; response: p = 0.210; feedback: p = 0.737, Figure 2d, purple, Table S3a). We next examined whether clusters demonstrated risk sensitivity, as would be expected if PCC sites are recruited in support of decisional processes. While we did not find evidence for risk sensitivity during the onset period in either task for type 1 sites (value-based, onset: p = 0.958; memory-based, onset: p = 0.476, Table S3b, Figure 2e), during the period following decision button-press responses, BBG activity during both tasks was significantly elevated for risky compared to safe trials (value-based, response: p = 0.002; memory-based, response: p = 0.038; Figure 2e). For the memory-based DM task, this risky-safe difference extended into the feedback presentation period, but not for the value-based DM task (memory-based feedback: p = 0.046; value-based, feedback: p = 0.211, Figure 2e, Table S3b). Conversely, type 2 sites did not demonstrate risk sensitivity during any time periods from either task (value-based, onset: p = 0.331; response: p = 0.874; feedback: p = 0.848; memory-based, onset: p = 0.093; response: p = 0.137; feedback: p = 0.246, Figure 2e, purple, Table S3b).

Beyond risk sensitivity, previous work has linked PCC activity to subjective value processing (Bartra et al., 2013; Kable & Glimcher, 2007; Lakhani et al., 2026). Therefore, we next examined whether PCC BBG responses during the value-based DM tracked subjective value (see methods). We found that type 1 BBG responses during the response period were related to subjective value, but not during the onset or feedback periods (value-based, onset: p = 0.170, response: p = 0.002, feedback: p = 0.053, Table S3c). Type 2 BBG responses were not modulated in relation to subjective value during any time period (onset: p = 0.855; response: p = 0.776; feedback: p = 0.608, Table S3c). These observations suggest risk sensitivity and value processing in PCC are largely restricted to type 1 sites, and temporally specific to post-decision phases, rather than during a pre-decision period.

#### Across-task similarity of PCC LFP response types

Next, we asked whether sites demonstrated similar engagement across the two DM tasks by directly testing whether the degree of BBG activity during value-based decisions was predicted by activity during memory-based decisions. For type 1 sites, responses were correlated during the initial onset and response periods, but not during feedback (across-task, onset: p < 0.001; response: p < 0.001; feedback: p = 0.083, Figure 2f, Table S3d). This pattern of correlations mirrors the risk sensitivity results, whereby risk-modulation was congruent across both tasks during the onset and response periods (Figure 2e), but incongruent during the feedback period. For type 2 sites, we did not observe correlated responses at any time period (across-task, onset: p = 0.572; response: p = 0.165; feedback: p = 0.216, Figure 2f, Table S3d), reflecting the low overall recruitment of these sites across both tasks.

### Distinct PCC single-unit response profiles during decision-making

We examined PCC microwire recordings to determine whether single-unit activity would be characterized by a homogenous, sustained pattern, resembling that observed in the population-level BBG LFP response, or instead distinct subsets differentially engaged across tasks and decision periods. To do so, we performed a similar data-driven clustering analysis as we did for PCC LFPs. For isolated units, baseline-corrected individual firing rate time series were extracted, concatenated across trial time periods, and averaged across trials within each block. A pairwise dissimilarity matrix was constructed between baseline-corrected firing rate time series, and hierarchical clustering was applied to identify distinct unit response types (Figure 3b, see methods for more details). Across tasks, we did not find consistent convergence for cluster assignments, and as such, clustering and subsequent analyses were performed separately for the value- and memory-based DM tasks.

**Figure 3.**
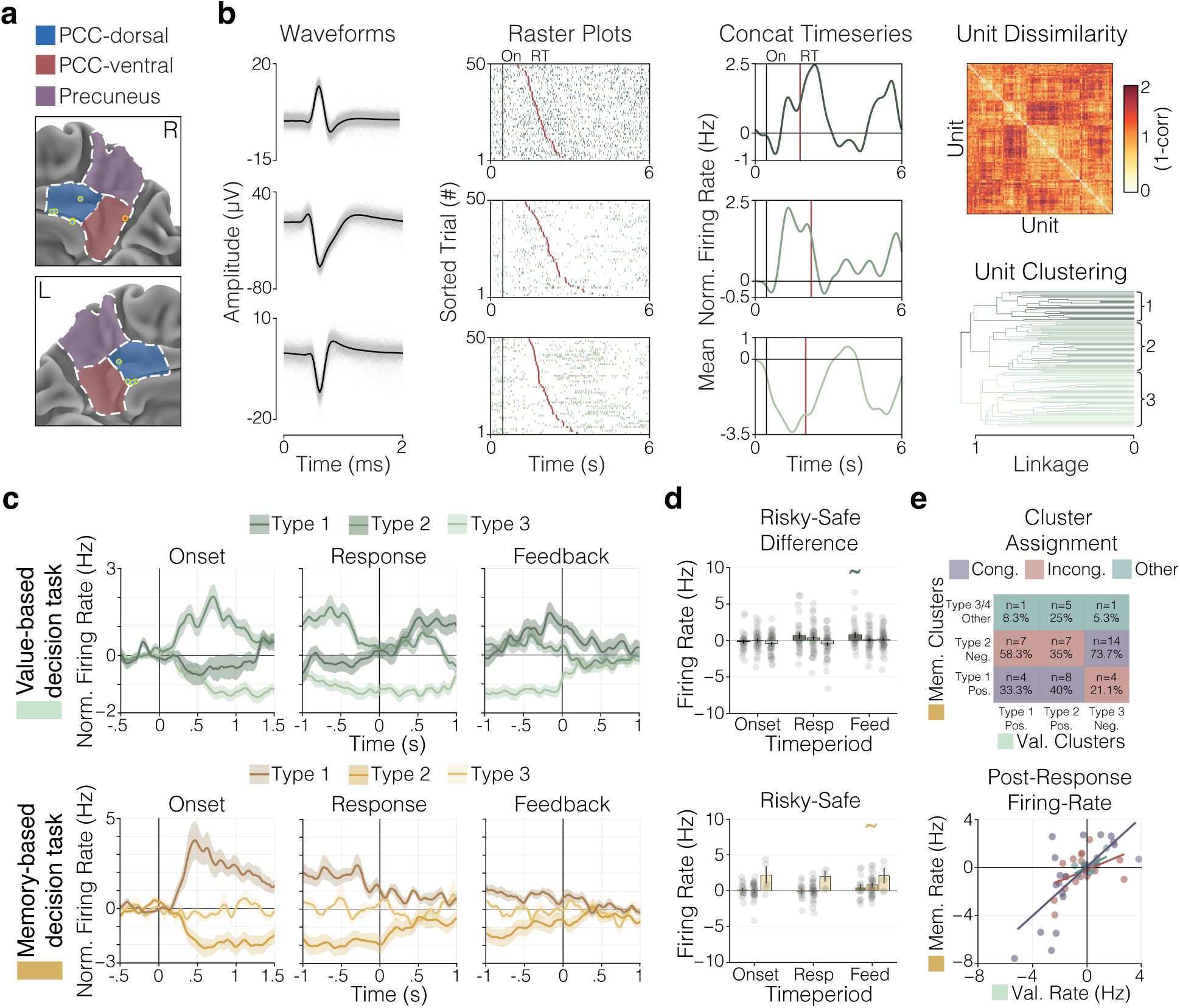
PCC single-unit responses and clustering of response types during decision-making. **a)** Focused view of MPC, highlighting anatomy and microwire recording site locations (blue = dorsal PCC, red = ventral PCC, purple = precuneus). Medial electrodes from probes containing microwires are indicated with yellow outlines (microwire sites n = 10). **b)** Single-unit data were analyzed following the clustering analysis performed for LFP data (Figure 2). For three example units from one value-based task block (from left to right): i) spike waveforms, ii) raster plot of spike timing aligned to trial onset and sorted by ascending RT (red), and iii) mean instantaneous firing rate time series constructed by concatenating across trial epochs, aligned to trial onset with mean RT indicated (red). iv) For all PCC units identified during the value-based task block (units n = 110), a pairwise dissimilarity matrix (1-r) of concatenated unit time series (top right) was used for hierarchical clustering (bottom right), resulting in three PCC unit types. **c)** Group-mean firing rate time series, normalized to the pre-trial baseline, for unit types identified during either the value-based (top) or memory-based (bottom) tasks. Responses are aligned to trial onset (left), decision button-press response (middle), and feedback presentation (right). **d)** The mean difference in firing rate for risky minus safe trials during the 500 ms period following trial onset, response, and feedback for unit types. **e)** Consistency of response profiles for units identified in both tasks. Classification agreement matrix showing the count and column-wise proportion of units with matching (congruent, purple), inverted (incongruent, red), or other (green) response profiles between the value-based (columns) and memory-based (rows) tasks (top). Group-mean post-response firing rate (500 ms) was strongly positively correlated across tasks for congruent units (purple), but no such relationship was observed for incongruent (red) or other (green) units (bottom).

#### Clustering reveals distinct PCC single-unit response types

For the value-based DM task, we identified three unit cluster types, and visualized their response across trial time periods (Figure 3c-top). Type 1 firing rates did not exceed baseline within the onset, response, and feedback time periods (value-based, onset: p = 0.139; response: p = 0.196; feedback: p = 0.084, Figure 3c-top, Table S4a). However, visual inspection of the type 1 time series revealed a consistent peak centered on feedback presentation, rather than the window taken after feedback presentation, during which firing rate was significantly elevated (value-based, feedback-centered: p = 0.034, Table S4a). Type 2 units demonstrated a different response pattern, only having increased activity during the trial onset period, but remaining at baseline for the other trial periods (value-based, onset: p = 0.032; response: p = 0.260; feedback: p = 0.587, Figure 3c-top, Table S4a). In contrast, type 3 units demonstrated sustained deactivation following trial onset and response periods, and returned to baseline after feedback presentation (value-based, onset: p = 0.038; response: p = 0.004; feedback: p = 0.271, Figure 3c-top, Table S4a). Risk sensitivity was observed for type 1 units during the feedback presentation period, though this effect did not remain statistically significant following multiple comparison correction (value-based, feedback: p = 0.035, FWER-corrected p = 0.057, Figure 3d-top, Table S4b). Interestingly, this result suggests that unit clustering distinguishes between units that separately underlie the early and late components of the sustained cluster-type 1 LFP response.

For the memory-based DM task, four types of unit response profiles were identified and characterized (Figure 3c-bottom), however, the fourth cluster was excluded from subsequent analyses due to insufficient membership (only 2 units). Type 1 memory-task units had increased firing rates following trial onset but not during the response or feedback periods (memory-based, onset: p = 0.035; response: p = 0.101; feedback: p = 0.129, Figure 3c-bottom, Table S4a), mirroring the response profile of type 2 value-task units. Type 2 memory-task units demonstrated a decrease in firing rate after trial onset but baseline firing rates during the response and feedback periods (memory-based, onset: p = 0.029; response: p = 0.272; feedback: p = 0.938, Figure 3c-bottom, Table S4a). Firing rate responses from type 3 memory-task units did not deviate from baseline at any timepoint (memory-based, onset: p = 0.457; response: p = 0.615; feedback: p = 0.574, Figure 3c-bottom, Table S4a). Only type 2 memory-task units demonstrated a degree of risk sensitivity, which was restricted to the feedback period and driven by attenuated deactivation for risky compared to safe trials, though this effect was not significant when corrected for multiple comparisons (memory-based, feedback: p = 0.022, corrected-FWER p = 0.157, Figure 3d-bottom, Table S4b).

#### Across-task similarity of PCC single-unit response types

We next examined unit generalizability across the value- and memory-based DM tasks (n = 51; Figure 3e). We first characterized the proportion of units where across-task cluster profiles were matched (congruent; e.g. positive/positive), inverted (incongruent; e.g. positive/negative), or mixed (other; e.g. positive/non-responsive). Approximately half of all units (51%, 26/51) demonstrated task congruent profiles, one-third were incongruent (35%, 18/51), and the remainder (14%, 7/51) were classified as ‘other’ (Figure 3e). We next quantified whether value-based response magnitude was predictive of memory-based response magnitude for the congruent, incongruent, and other units. For congruent units, value- and memory-based responses were correlated following the decision button-press response and feedback presentation, but not onset period (across-task, onset: p = 0.519; response: p < 0.001; feedback: p = 0.022, Figure 3e, Table S4c). Incongruent units demonstrated negative correlations after trial onset, no systematic relationship after response, and a positive correlation after feedback presentation (across-task, onset: p = 0.016; response: p = 0.166; feedback: p = 0.010, Figure 3e, Table S4c), while other units did not show systematic correlations at any timepoints (across-task, onset: p = 0.554; response: p = 0.471; feedback: p = 0.433, Figure 3e, S4c). These results demonstrate that while some units show domain-general response profiles across both DM tasks, mirroring the pattern observed from PCC LFPs, a separate subset of units show selectivity dependent on the type of information driving decisions. Altogether, these data suggest that the observed sustained LFP response in PCC reflects the aggregation of distinct single-unit subpopulations, each engaged at different stages of the decision process, revealing a heterogeneous functional organization underlying population-level responses in PCC.

### Hippocampal LFP responses during decision-making

The contrasting of value- and memory-based tasks used in the current study also offers an opportunity to examine broader mnemonic network engagement during decisional processes. Given a growing literature implicating the hippocampus in both decision contexts (Bakkour et al., 2019; Biderman et al., 2020; Lopez-Persem et al., 2020), we next assessed hippocampal LFP and single-unit responses across value- and memory-based tasks (Figure 4a). The hippocampal sample comprised both PCC participants with concurrent hippocampal coverage and additional participants with only hippocampal recording sites (see supplementary Table S1 for details). Probes were labeled as anterior or posterior hippocampus based on their position relative to the uncal apex (Poppenk et al., 2013, see methods for details), consistent with the distinct structural, functional, and connectivity profiles observed along the hippocampal long-axis (Barnett et al., 2021; Strange et al., 2014; Zheng et al., 2021). This division is particularly relevant given recent evidence for preferential links between posterior hippocampus to dPCC and the anterior hippocampus to vPCC (Angeli et al., 2025; Koslov et al., 2024).

**Figure 4.**
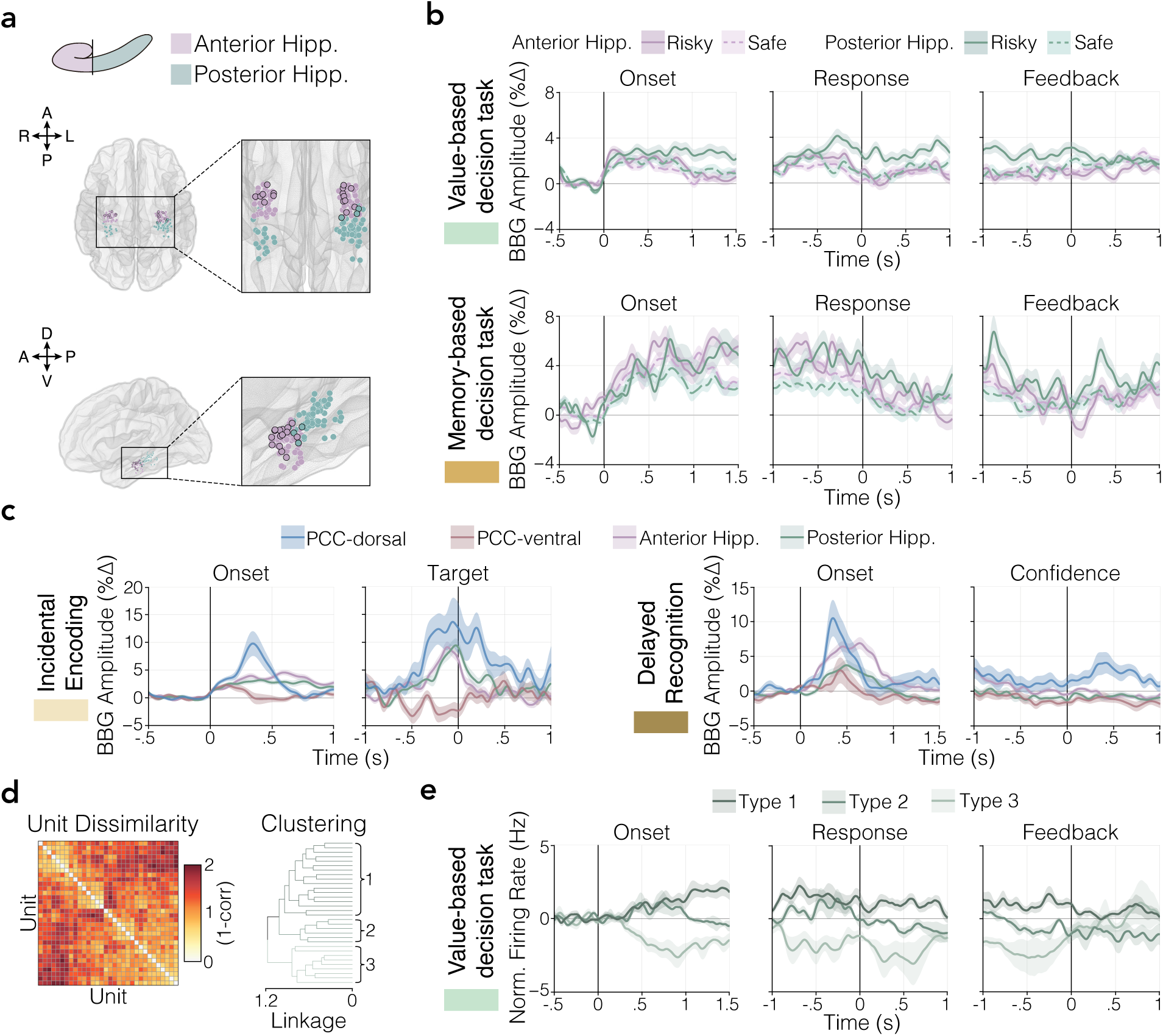
Hippocampal LFP responses during decision-making, encoding and delayed recognition tasks, compared to PCC, and hippocampal single-unit responses during value-based decision-making. **a)** Schematic illustrating the anatomical criterion used to label hippocampal recording sites as anterior (anterior hipp., purple) or posterior (posterior hipp., green), based on position relative to the uncal apex (top). Wire mesh rendering of MNI152 cortex showing the normalized hippocampal recording site locations across participants (bottom; macro electrodes n = 142). Insets highlight the distribution of recording sites across the hippocampus, with distal electrodes of probes containing microwires outlined in black (microwire sites n = 19). **b)** Group-mean BBG (70-150 Hz, percent signal change from pre-trial baseline) LFP responses on risky (solid lines) and safe (dashed lines) trials during the value-based (top) and memory-based (bottom) tasks from the anterior hipp. (purple) and posterior hipp. (green). Responses are aligned to trial onset (left), decision button-press response (middle), and feedback presentation (right). **c)** Group-mean BBG responses during the incidental 1-back encoding task aligned to trial onset (far-left) and 1-back target responses (central-left) from electrodes in the dPCC (blue), vPCC (red), anterior hipp. (purple), and posterior hipp. (green). Group-mean BBG LFP responses during the delayed recognition task aligned to trial onset (center-right) and confidence onset (far-right) from electrodes in the dPCC (blue), vPCC (red), anterior hipp. (purple), and posterior hipp. (green). **d)** Hippocampal single-unit data were analyzed following the clustering analysis performed for PCC units (Figure 3b). For units (n = 32) isolated from value-based task microwire recordings, a pairwise dissimilarity matrix (1-r) of concatenated unit timeseries (left) was used for hierarchical clustering (bottom right), resulting in three hippocampal unit types. **e)** Mean firing rate timeseries (normalized to the pre-trial baseline) for the three hippocampal unit types aligned to trial onset (left), decision button-press response (middle), and feedback presentation (right).

We first analyzed LFPs, focusing on BBG activity, from anterior and posterior hippocampal recording sites (total n = 142, anterior n = 83, posterior n = 59). A data-driven clustering approach for hippocampal LFPs failed to yield at least two clusters of sufficient size, and as such, anatomical labels were used for subsequent task analyses. For the value-based DM task (Figure 4b-top), both anterior and posterior hippocampal sites demonstrated BBG amplitude significantly above baseline during the trial onset and feedback time periods, however only posterior sites were above baseline following responses (value-based, anterior, onset: p < 0.001; response: p = 0.164; feedback: p = 0.003; posterior, onset: p < 0.001; response: p = 0.019; feedback: p < 0.001, Figure 4b-top, Table S5a). Interestingly, response magnitude from posterior sites was significantly greater than anterior sites during all time periods (value-based, anterior vs. posterior, onset: p = 0.009; response: p = 0.003; feedback: p < 0.001, Table S5b). Furthermore, only posterior hippocampus sites demonstrated risk-sensitivity, with greater BBG following responses for risky compared to safe trials (value-based, risk-modulation response: p = 0.004, Table S5c). For the memory-based DM task, both anterior and posterior hippocampus demonstrated above baseline BBG responses during all time periods (memory-based, anterior, onset: p < 0.001; response: p = 0.006; feedback: p = 0.018; posterior, onset: p < 0.001; response: p = 0.027; feedback: p = 0.005, Figure 4b-bottom, Table S5a), however, there was no difference in response magnitude across the two subregions (memory-based, anterior vs. posterior, onset: p = 0.234; response: p = 0.470; feedback: p = 0.109, Table S5b). Like the value task, only posterior hippocampus demonstrated risk sensitivity during the memory-based DM task, with greater responses to risky than safe trials after feedback (memory-based, risk-modulation, feedback: p = 0.029, Table S5c). Overall, anterior and posterior hippocampal subregions showed differential engagement across DM tasks. Notably, the response profile of posterior hippocampus, marked by sustained engagement and risk sensitivity across both tasks, mirrors that of the mainly dorsal PCC cluster 1, consistent with a preferential link between the posterior hippocampus and dPCC.

### Across-region LFP responses during encoding

To further characterize PCC and hippocampal recruitment during memory-based decisional processes, we next looked at BBG amplitude during the incidental encoding 1-back task for dPCC, vPCC, anterior hippocampus, and posterior hippocampus recording sites (Figure 4c). Following trial onset, significant increases in BBG amplitude were observed from dPCC, anterior hippocampus, and posterior hippocampus, but not vPCC recording sites (trial onset, dPCC: p = 0.003; vPCC: p = 0.176; anterior: p < 0.001; posterior: p < 0.001, Table S6a). Interestingly, dPCC sites demonstrated a significantly larger BBG amplitude increase after trial onset compared to other sites (vs. dPCC, vPCC: p < 0.001; anterior: p = 0.003; posterior: p < 0.001, Table S6a). Similarly, during 1-back target responses, BBG activity was significantly elevated for dPCC, anterior hippocampus, and posterior hippocampus, but not for vPCC recording sites (target onset, dPCC: p = 0.005; vPCC: p = 0.639; anterior: p = 0.008; posterior: p < 0.001, Table S6b). Again, dPCC demonstrated significantly larger amplitude of BBG responses compared to other regions (vs. dPCC, vPCC: p < 0.001; anterior: p < 0.001; posterior: p = 0.006, Table S6b). Together, these results reinforce the functional distinction between dorsal and ventral PCC, while also again demonstrating similarities in timing, but differences in magnitude, of dPCC and hippocampal subregional recruitment during memory task performance.

### Across-region LFP responses during delayed recognition

Participants also completed a delayed recognition task, with distinct item-recognition trial onset and confidence response periods, from which we examined PCC and hippocampal LFP responses. Following trial onset, BBG responses were significantly elevated for dPCC, anterior hippocampus, and posterior hippocampus, but not vPCC (trial onset, dPCC: p = 0.004; vPCC: p = 0.272; anterior: p < 0.001; posterior: p = 0.012, Table S7a). Trial onset response amplitudes were similar between sites in the dPCC and anterior hippocampus (vs. dPCC, anterior: p = 0.309, Table S7a), but greater for dPCC compared to vPCC and posterior hippocampus (vs. dPCC, vPCC: p = 0.001; posterior: p = 0.004, Table S7a). Interestingly, during the confidence-response period, elevated BBG responses were observed only for dPCC (confidence onset, dPCC: p = 0.032; vPCC: p = 0.311; anterior: p = 0.547; posterior: p = 0.066, Table S7b), and dPCC responses were statistically greater than for all other sites (vs. dPCC, vPCC: p < 0.001; anterior: p < 0.001; posterior: p < 0.001, Table S7b). While the convergence of dPCC and hippocampal recruitment during recognition-memory decisions following trial onset further supports a role for the dPCC in memory-based decision-making, the unique engagement of dPCC during confidence decisions also points to a broader role for the region in domain general decisional processing.

### Distinct hippocampal single-unit response during value-based decision-making

Finally, we characterized response profiles of hippocampal units isolated from the value-based DM task, using the same data-driven clustering approach that was applied to PCC units (see methods for details). As the region was a secondary area of focus, there were fewer resulting single-units from the hippocampus, and subsequently, sufficient data (units n = 32) for analysis were only obtained during the value-based DM task. Three distinct response profiles were identified for hippocampal units (Figure 4d) and characterized across time periods (Figure 4e). Type 1 units showed a modest increase in firing rate only following trial onset, but did not statistically differ from baseline during any task time period (value-based, onset: p = 0.297; response: p = 0.234; feedback: p = 0.299, Table S8a). Firing rates for type 2 units were of similarly low magnitude (value-based, onset: p = 0.422; response: p = 0.137; feedback: p = 0.362, Table S8a), only increasing above baseline during a period centered on feedback presentation (feedback-centered: p = 0.046, Table S8a). This pattern, with increased activity centered around the time of feedback presentation, mirrors that of PCC type 1 units. Hippocampal type 3 units had significantly suppressed firing rates during the onset and response time periods, returning to baseline following feedback (value-based, onset: p = 0.007; response: p = 0.026; feedback: p = 0.779, Table S8a), mirroring PCC type 3 units. Unlike PCC, no hippocampal unit type demonstrated risk sensitivity (Table S8b). Despite fewer available units, response profiles from hippocampal units broadly echoed those observed from PCC units, suggesting concurrent recruitment during value-based decision-making. More generally, single-unit results indicate that sustained population-level LFPs in both PCC and hippocampus reflect the aggregation of distinct single-unit subpopulations, each preferentially engaged or inhibited at different periods of decision-making.

## Discussion

To better understand how the human PCC contributes to decision-making, we obtained multiscale LFP and single-unit recordings while participants performed matched value- and memory-based decision tasks. Anatomically, analysis of LFP responses, focused on BBG activity, revealed a clear dissociation between PCC subregions, whereby dPCC, but not vPCC, was engaged across both decision-making tasks. Physiologically, LFP responses in dPCC showed a similar sustained temporal profile throughout decision-making in both tasks. However, it was only in the post-decision period that LFP responses correlated with choice risk and subjective value. In contrast to the population-level LFP responses, single-unit activity was more heterogeneous and temporally circumscribed, clustering into differing temporal profiles of task response. Specifically, dPCC unit firing was either suppressed throughout the decision trial, or selectively increased in the pre- or post-decision period. Unlike LFP responses, single-unit activity profiles were only consistent across tasks for a subset of units. To situate PCC activity within a broader memory network, we examined LFP responses in the hippocampus, observing temporal profiles similar to those of dPCC. Together, these anatomical and physiological features of human PCC responses during decision-making provide a compelling link between prior work in NHP electrophysiology and human neuroimaging, further promoting PCC function as a convergence of both executive and mnemonic processing.

Our electrophysiological findings revealed a striking anatomical dissociation of responses between human dorsal and ventral PCC across decision-making tasks. This subregional dissociation is consistent with a substantial body of human neuroimaging research that together sheds light on the putative functional neuroanatomy of PCC (Foster et al., 2023; Foster & Koslov, 2025). For example, in studies examining differing tasks of episodic memory retrieval, a consistent dissociation has been observed whereby item-recognition decisions are uniquely associated with dPCC, while autobiographical retrieval is predominantly associated with vPCC activity (Chen et al., 2017; McDermott et al., 2009). Convergent with these findings, human neuroimaging studies also implicate dPCC in memory-guided decisions that require temporally extended information or long-term memory (Law et al., 2023; Rosen et al., 2018; Visalli et al., 2019), as well as in value-based decision making (Engelmann & Tamir, 2009; Nieuwenhuis et al., 2005), together pointing to the dPCC as a convergence site for evaluative and mnemonic processes. Furthermore, prior work has shown that while dPCC is engaged during memory-based decisions, the vPCC is more associated with the representation of related episodic and semantic content of such decisions (Binder et al., 2009; Elman et al., 2012, 2013; Kim, 2021; Koslov et al., 2024). The differing profiles of dPCC and vPCC responses align with recent precision functional neuroimaging studies, which link dPCC to frontoparietal control and salience networks and associate vPCC with the broader DMN (Angeli et al., 2025; Du et al., 2024; Gordon, Laumann, Gilmore, et al., 2017; Kwon et al., 2025). This pattern is also consistent with earlier reports of subregional differences in cytoarchitecture and structural connectivity between these subregions (Foster et al., 2023; Greicius et al., 2009; Kobayashi & Amaral, 2000; Vogt et al., 1995; Vogt & Palomero-Gallagher, 2012). Together, this refined functional neuroanatomy helps reconcile prior challenges when interpreting NHP PCC electrophysiology in light of human neuroimaging (Pearson et al., 2011). Because NHP experiments were almost exclusively performed in dPCC, focusing on single-unit responses to economic decision behavior, prior efforts to relate these data to the earlier human emphasis on the DMN and episodic memory presented an apparent tension that resolves through clarifying the subregional homology between species. Through this anatomical insight, progress can also be made on existing open questions regarding the timing of PCC engagement during decision-making.

In addition to spatial dissociation, multi-scale electrophysiological recordings offer insight into the physiology of human PCC engagement during decision-making. For example, prior human neuroimaging studies have differed in linking dPCC activity to either pre- or post-decision periods (e.g. Bartra et al., 2013; Clithero & Rangel, 2014; Oldham et al., 2018), a timing question well suited to complementary methods, such as invasive electrophysiology, which provide fine-grained temporal resolution. Across both the value- and memory-based decision-making tasks in the current study, dPCC BBG LFP responses were sustained from trial onset until shortly after feedback presentation, with risk-sensitivity observed only during the post-decision period. As a temporal correlate of fMRI BOLD (Hermes et al., 2012; Mukamel et al., 2005; Nir et al., 2007), BBG allows these findings to inform prior neuroimaging literature, suggesting that studies reporting either pre- or post-decision dPCC activity likely captured different facets of this sustained engagement. Additionally, given that BBG is also a temporal correlate of local population spiking activity (Lei et al., 2026; Mukamel et al., 2005; Nir et al., 2007; Ray & Maunsell, 2011), the single-unit data suggest the sustained LFP response reflects an aggregation of neurons engaged during different timepoints throughout task trials. Specifically, we observed separate clusters of units that increased firing during either the pre-decision or post-decision period. The heterogeneous timing of unit responses initially appears at odds with prior NHP work, which has predominantly reported engagement of single-unit activity during the post-decision period (Hayden et al., 2008; Heilbronner et al., 2011; Heilbronner & Platt, 2013; Li et al., 2019). However, this discrepancy results in part from differences in study goals and resulting analytical approaches, in that many NHP studies focused on isolating specific decision response types, a subset of all isolated units, for in-depth analysis. By clustering all isolated single-units, we identified units with pre-decision increases in firing. Indeed, a subset of prior NHP studies that reported broader unit samples or population-level responses also observed evidence for dPCC engagement during both pre- and post-decision periods (Fine et al., 2023; Heilbronner & Platt, 2013; McCoy et al., 2003; McCoy & Platt, 2005). In addition, our findings and prior NHP studies (Hayden et al., 2009, 2010) both identified a subset of cells whose firing is consistently suppressed throughout task trials. Given that PCC was historically noted as a region associated with task-related deactivation (Fox et al., 2005; Shulman et al., 1997), it is likely important to consider this full complement of unit response types for understanding the region’s underlying neural computations. Finally, in line with prior NHP findings, we identified a subset of units that demonstrated a degree of post-decision risk-sensitivity, though across the subset this effect was small and did not survive multiple comparison correction. Altogether, these physiological aspects of our findings address important questions about the timing of PCC subregional recruitment during decision-making, and suggest a compelling alignment of findings across species and scales of measurement.

In addition to the above advances, there are important limitations of our study to consider for future investigations into PCC function. Overall, our findings draw focus to dPCC rather than vPCC, a distinction likely influenced by the tasks employed. This is consistent with vPCC being more associated with episodic and contextual details of behavior (Andrews-Hanna et al., 2014; Bird et al., 2015), cognitive domains that future invasive studies could target directly. Such work could also employ task paradigms in which decisions require a more naturalistic use of sensory or mnemonic information, enabling study of how the subregions interact (Gilmore et al., 2021; Lee & Chen, 2022; Masís-Obando et al., 2022; Su et al., 2025). In considering future investigations, it is important to note the correlational nature of our observations, and the open question regarding the causal role of PCC in decision-making. This is a particularly notable gap, as unlike other brain structures, there is little substantive neuropsychological literature documenting the behavioral consequences of PCC damage. Ultimately, such causal evidence is particularly important for elucidating the role of PCC in disease. Finally, the single-unit heterogeneity observed here reflects a sparse sample of a much larger neural population. Recording from greater numbers of cells, and analyzing their collective population-level structure, would help elucidate the coding mechanisms of PCC subregions and further link findings across species and measurement scales.

In summary, the results from the current study reconcile findings of PCC engagement across separate research domains and species. An important part of resolving previous disparities comes from considering the functional distinctions between dPCC and vPCC subregions, instead of treating PCC as a unitary region. Even when considering dPCC alone, no single account, ranging from a DMN-centered role in episodic memory to a role in economic valuation and outcome monitoring, fully explains the diversity of processes ascribed to the region, instead pointing to a domain-general or higher-order function. Consistent with this domain-general account, we observed similar dPCC responses across value- and memory-based decisions, as well as engagement during incidental encoding, delayed recognition, and confidence judgments. This domain-general account of dPCC function, however, does not preclude important dPCC contributions to memory-related processes. Prior work establishing dPCC as a connector hub between control and default networks (Gordon et al., 2018), the region’s associations with item-recognition and memory-guided attention (Isenburg et al., 2023; Kim, 2013; Rosen et al., 2018; Wagner et al., 2005), and the close correspondence with hippocampal responses observed here together support a role for dPCC at the intersection of decision-making and memory. Beyond these task-specific associations, however, the region’s contributions may be better revealed at a higher-order level, operating over longer timescales — consistent with the view that PCC sits near the apex of cortical processing hierarchies, integrating information across extended periods (Dohmatob et al., 2020; Foster & Koslov, 2025). On this account, dPCC’s sustained, domain-general engagement may reflect not the momentary computation of a decision, but the ongoing integration of accumulated context and prior experience with incoming information — a process suited to representing past and anticipated behavioral scenarios rather than isolated choices. Although testing these claims will require targeted study, our findings provide a critical alignment, previously missing from the literature, across cognitive domains, species, and measurement scales, helping to orient the field to a compelling path for understanding this historically enigmatic region.

## Methods

### Human participants

Intracranial recordings were obtained from 38 human participants, reflecting PCC and/or hippocampal recordings (19 F, 19 M; mean age = 37.1 yrs, age range: 21-68 yrs; see Table S1) undergoing invasive monitoring for refractory epilepsy at the University of Pennsylvania (Philadelphia, PA, USA), Medical College of Wisconsin (Milwaukee, WI, USA), or Baylor St. Luke’s Hospital (Houston, TX, USA). Participant information and recording site details are in table S1. Participants with epileptic foci localized to the posterior cingulate cortex were excluded. All experimental procedures were approved by the Institutional Review Boards at the University of Pennsylvania Perelman School of Medicine (#: 821778), Baylor College of Medicine (#: H-18112), and Medical College of Wisconsin (#: 44904), with patients providing verbal and written consent.

### Experimental design

Participants performed a value-based decision-making task and a three-part memory-based encoding, decision-making and recognition paradigm across multiple recording sessions/days (see Figure 1b). Task timing varied with participant availability during clinical monitoring, and as such, not every participant completed all task components (see Table S1 for details). Across participants, however, task structure and ordering remained consistent. Participants first completed the value-based decision-making (DM) task, followed by an incidental encoding 1-back task, a memory-based DM task, and after a delay of approximately 48 hrs, a delayed recognition task. All tasks were presented on a monitor placed at a comfortable distance and position for participants, and run using MATLAB (R2023a, MathWorks, MA, USA) and Psychtoolbox (3.0.19; Brainard, 1997).

### Value-based decision-making task

The structure of the value-based DM task followed established designs used in previous studies of risky decision-making in both humans (e.g. Glickman et al., 2019; Jung et al., 2018; Saez et al., 2018) and non-human primates (Fine et al., 2023; Heilbronner et al., 2011). On each trial of the value-based DM task, participants chose between a safe and risky option presented simultaneously on the screen (see Figure 1c-top), indicating their choice by a right or left keypad button-press. The safe option always offered a $10 reward with 100% probability. The risky option offered a higher reward ($12, $15, $20, $25, or $30) at a probabilistic chance (20-90%). Reward probabilities were conveyed via spin wheels divided into 10 color-coded slices (green = win, red = loss), with the number of green slices explicitly indicating the probability of reward. Reward value was displayed numerically above each wheel, and losses returned no reward ($0). Value-probability pairings were randomly drawn without replacement from the factorial combination of all value and probability levels (40 total). To further encourage choice deliberation, 10 of the pairings with expected values (EV = value * probability) approximating the safe option EV of 10 were repeated during each block. The left/right location of the safe and risky options was counterbalanced across trials. Participants had up to 4500 ms to make their decision, with non-responses prompting a message to ‘try to respond faster on the next trial’. Immediately following the button-press choice decision, a yellow box appeared around the chosen option and the wheel’s needle rotated around clockwise for between 500 – 1200 ms, with its landing position indicating trial outcome. Feedback was then presented in the form of both the yellow box around the chosen option changing colors (green = win; red = loss) and text (‘win’ for wins or ‘unlucky’ for losses) appearing on the screen for 1000 ms. Feedback was followed by an ITI period of 500 – 1000 ms. Participants completed 50 trials per block (∼4-6 minutes each) with most participants finishing 2 consecutive blocks in an initial session. In a subsequent session, participants performed an additional 1-2 blocks, after completing the incidental encoding and memory-based DM tasks. Due to the variability of participant availability, total block completion varied across individuals (see Table S1 for details).

### Memory paradigm part 1: incidental encoding 1-back task

The memory-based DM task was designed to match the structure and timing of the value-based DM task, differing only in the need for mnemonic information to support decision-making. To do so, we had participants complete a three-stage memory paradigm which included an initial encoding 1-back task, a memory-based decision-making task, and a delayed recognition task (Figure 1b). The encoding and memory-based DM tasks were completed consecutively and performed after the initial blocks of the value-based DM task.

For the incidental encoding 1-back task (Figure 1e), grayscale images of faces, houses, and vehicles (Stigliani et al., 2015) were presented sequentially for 1000 ms each, with participants instructed to make a keypad button-press response upon detecting an immediate repeat (1-back). We sought to manipulate the memory encoding strength of stimuli that would be subsequently used for the memory-based DM task by presenting stimuli either 1, 2, 3, or 5 times during encoding, with more frequently encoded stimuli expected to be more readily remembered on average. An equal number of faces, houses, and vehicles were used (54 unique images), with repetition frequency levels equally distributed across the encoding task. An additional 15 images served as 1-back targets (presented twice consecutively) and 15 scrambled background images were randomly interspersed within the encoding task, with neither set of images used in subsequent tasks. The encoding task included 180 total trials and took approximately 7.5 minutes.

### Memory paradigm part 2: memory-based decision-making task

Immediately following the encoding task, participants completed the memory-based DM task (Figure 1c-bottom), which was matched in timing and structure to the value-based DM task, differing only in the inclusion of mnemonic information to drive decisions. Participants were told that points would be earned during the memory-based DM task by selecting stimuli that had been seen at any previous point in the experiment (either during encoding or the current DM task). On each trial, participants chose between a safe and risky option presented simultaneously on the screen (see Figure 1c-bottom), indicating their choice by a right or left keypad button-press. The safe option always paired a $10 reward with a frequently seen stimulus (i.e. 100% probability of reward). A frequently seen stimulus was always used as the safe option, and safe-stimuli were repeatedly used throughout the DM task, such that participants quickly learned that the $10 option always returned a reward. The risky option offered a higher reward ($20, $25, or $30), but was paired with an image that was equally likely to be an infrequently encoded (‘old’) stimulus or a novel (‘new’) stimulus not seen during encoding. Response window, feedback structure, and ITI were identical to the value-based DM task, with the exception that after a button-press response a semi-transparent circle timer rotated clockwise over the chosen stimulus for 500 – 1200 ms in place of a spinning wheel. Participants completed two consecutive memory-based DM task blocks of 45 trials each (∼4-6 minutes per block).

### Memory paradigm part 3: delayed recognition task

After a delay period (mean = 47.4 hours, for details see Table S1), participants completed a delayed recognition task (Figure 1f). On each trial, a single stimulus was presented, and participants had up to 7000 ms to make a seen/unseen (“old”/“new”) decision using a left or right keypad button-press. A black box would appear around the chosen option immediately after selection, followed by a 250-750 ms delay. Every recognition decision trial was followed by a 5-point scale prompt where participants had up to 7000 ms to use keypad button-presses to indicate decision confidence. The confidence response scale ranged from “not sure” to “completely sure” with a midpoint of “maybe sure”. Participants were instructed to select “not sure” if they were completely guessing and “completely sure” if they were 100% certain about their memory decision. No feedback was provided during this task. In order to limit participant fatigue, the delayed recognition task was split into two blocks, one of 75 trials and the other of 78 trials (∼7-15 minutes per block), which were performed consecutively with a short break in between. Each block was comprised of 27 unseen/’new’ stimuli and 27 seen/‘old’ stimuli from the initial encoding period. Also included were 21 stimuli (24 for block 2) that had served as the ‘novel’-risky option during the memory-based DM task (considered as seen/‘old’ for the delayed recognition task), for additional exposure recency and repetition time comparison. Due to the variability of participant availability, total block completion varied across individuals (see Table S1 for details).

### Behavioral analyses

Behavior was analyzed from participants with PCC macro electrode recording sites. We used linear mixed-effect models, with a random intercept term for participant, and when applicable, a random slope term for participant. Value-based DM task (n = 20) performance was analyzed using mixed-effects logistic regression, comparing trial-wise likelihood of selecting the risky option as a function of the risky option’s expected value. The group model included random intercept and slope terms for participant. For the memory-based DM task (n = 15), behavior was analyzed by comparing the likelihood of choosing the risky option as a function of the number of times the risky option had been repeated during the incidental encoding 1-back task (0, 1, 2, or 3 times). The mixed-effects linear regression group model included a random intercept term for participant. For the delayed recognition task, recognition memory was analyzed by comparing d-prime across five stimulus repetition levels, using mixed-effects linear regression with a random intercept term for participant. Stimulus levels for seen/’old’ stimuli were: 1 (stimuli presented only as the unseen/‘novel’ option during the memory-based DM task), 2, 3, and 4 (stimuli presented 1,2, and 3 times during encoding and then once again as the ‘risky’ option during the memory-based DM task), or 5 (stimuli presented 5 times during encoding and then repeatedly as the ‘safe’ option during the memory-based DM task). For each repetition level, d-prime was calculated from the hit rate relative to the participant’s overall false alarm rate.

### Electrophysiological recordings

Recordings were performed using stereo-electroencephalography (sEEG) depth electrodes and Behnke-Fried macro-micro probes (Ad-Tech Medical Instruments Corp., WI, USA). Behnke-Fried electrodes included a bundle of 8 shielded microwires and 1 unshielded reference microwire (38-40 μm in diameter each, extending approximately 4mm from the distal tip of the probe). Microwire recordings were obtained from PCC (n = 10) and hippocampal (n = 19) locations. All sEEG trajectory decisions were made solely on the basis of clinical criteria.

Intracranial data were acquired at two electrophysiological scales: macro electrode LFP and microwire unit recordings. LFP data were acquired at a sampling rate of 2 kHz with a bandpass of 0.3 – 500 Hz (4^th^-order Butterworth filter) using a Blackrock Cerebus system (Penn / BCM) and Ripple Neuro (MCW). Initial recordings were referenced to a depth electrode contact within either skull or white matter, distant from pathological zones. Microwire data were simultaneously acquired at a sampling rate of 30 kHz using a Blackrock Cerebus system. Task event timing was tracked using a photodiode sensor (attached to the stimulus screen) synchronously recorded at 30 kHz. All subsequent processing was performed offline.

### Electrode localization

Electrode locations within PCC and hippocampus were identified in native subject space for each participant by co-registering a post-operative CT scan to a pre-operative T1 anatomical MRI using a combination of the YAEL (Your Advanced Electrode Localizer, Wang et al., 2023) module from RAVE (Reproducible Analysis and Visualization of iEEG software, Magnotti et al., 2020), FSL (FMRIB Software Library, Jenkinson et al., 2012), and ANTs (Advanced Normalization Tools, Avants et al., 2011). PCC was defined by boundaries at the marginal ramus of the cingulate sulcus, parieto-occipital sulcus, splenial sulcus, and corpus callosum (see Figure 1a). Dorsal and ventral subregions were demarcated using regional sulci along recent definitions (Foster et al., 2023; Willbrand et al., 2022). Hippocampal electrodes were manually verified to ensure that they were located within the hippocampus and to exclude neighboring regions. Electrodes were assigned to the anterior or posterior hippocampus based on position relative to the uncal apex as identified in each participant’s native-space anatomical image (Poppenk et al., 2013; Poppenk et al., 2020). For visualization purposes, all electrodes were normalized into MNI152 space and inspected for accuracy using RAVE and ANTs tools.

### Macro electrode signal processing and spectral decomposition of LFP activity

All macro electrode LFP signal processing was performed using custom MATLAB (v2021b, MathWorks, MA, USA) scripts. Raw signals were first inspected for line noise, recording artifacts, and interictal epileptic spikes. Electrodes with clear artifactual or persistent epileptic activity were excluded from further analysis. Next, signals were notch filtered (60Hz and harmonics) and bipolar re-referenced to an adjacent electrode. Re-referenced signals were then downsampled to 1 kHz and spectrally decomposed using Morlet wavelets (7 cycles), with center frequencies spaced linearly from 1 to 200 Hz in 1 Hz steps. For the incidental encoding 1-back task, value-based DM task, and memory-based DM task the instantaneous amplitude across spectral frequencies was converted to percent signal change by applying a pre-trial baseline correction (−501 to −1 ms for each trial). For the delayed recognition task, because ITIs were often shorter than 500ms, the mean activity from the entire task was used for the same baseline correction. The percent signal change time series was again inspected for interictal spiking activity and artifactual noise, and any trials where interictal spikes were visually observed were excluded from further analysis. Broadband gamma (BBG), defined as neural activity between 70 to 150 Hz, was extracted for each trial. In order to examine neural responses across separate decision-making stages while also accounting for variable trial durations, time series were aligned to trial onset, decision button-press response, and feedback presentation times.

### Micro electrode signal processing and spike sorting

Raw microwire signals were visually inspected and concatenated across blocks from the same recording session, ensuring confident identification of the same units across consecutive blocks and tasks. For recordings with evidence of line-noise or narrow-band environmental artifacts, adaptive notch filtering was applied following a data-driven noise removal procedure (Betancourt et al., 2026). Spike detection and sorting were then performed using WaveClus3 (Chaure et al., 2018) in MATLAB (v2019b). Signals were bandpass filtered between 300 to 3000 Hz, and an absolute voltage threshold for spike detection was set independently for each recording as 5 times the absolute median deviation. Putative spike waveforms were extracted around each threshold crossing, features derived using wavelet decomposition, and nonparametric clustering was applied to features. A semi-automated procedure was then used to optimize the number and size of clusters. Clusters were visually inspected, and those deemed to potentially contain single- or multi-units were retained for further analysis. A Duplicate Event Removal algorithm (DER, Dehnen et al., 2021) was applied to remove duplicate spikes identified as repeated across wires, bundles, and probes. Next, putative units were manually inspected for waveform consistency, canonical action potential morphology, and ISI refractory period violations (<3ms). Following this consensus verification process, 10 PCC and 9 hippocampal Behnke-Fried probes (from n = 10 and n = 9 participants, respectively) yielded units for subsequent analyses, with 110 PCC and 32 hippocampal units identified during the value-based DM task. However, single-unit yields from the memory-based DM task were limited due to fewer participants completing the task. As such, we were able to identify 52 PCC units from the memory-based DM task, but had insufficient hippocampal unit data for further analyses. Firing rates were computed for each trial, baseline corrected by subtracting the mean firing rate from the pre-trial baseline period (−501 to −1 ms), and aligned to trial onset, decision button-press response, and feedback presentation.

### Functional clustering of LFP and single-unit activity

Clustering analyses were performed in a similar fashion for PCC LFPs, PCC units, and hippocampal units. Preprocessed data (percent signal change for LFPs and baseline-corrected firing rate for units) were extracted for each trial. To combine data across trials of varying lengths and capture response dynamics in relation to the separate stages of decision-making, timeseries of fixed length were concatenated across four periods: trial onset (−500 to 1500 ms), decision button-press response (−1000 to 1000 ms), feedback presentation (−1000 to 1000 ms) and ITI (0 to 500 ms). Trial-wise concatenated time series were then averaged such that there was one time series per electrode or unit per block. Next, we sought to identify any potential similarities in patterns of responses across macro- or micro-recordings by performing unsupervised hierarchical clustering of the time series data. For clustering, all concatenated time series (example Figure 2c, left) from one block were used to calculate pairwise Pearson correlations, which were then inverted to generate a pairwise dissimilarity matrix (1-r; Figure 2c, center-left). Hierarchical clustering was performed on the dissimilarity matrix in R, and the optimal number of clusters was selected by evaluating the combination of within-cluster sum of squares, silhouette width peak, gap statistics, and PCA decomposition features of the dissimilarity matrix. For PCC LFPs, clustering was initially performed using the first value-based DM task block and compared to clustering using the second value-based DM task block and either the first or second memory-based DM task block. As qualitatively similar results, yielding 5 cluster types, were observed across clustering blocks, the clustering outputs from the first value-based DM task block were used for subsequent analyses. PCC unit clustering followed the same procedure, however, clustering assignments from the value- and memory-based DM tasks differed and were subsequently analyzed independently. Due to fewer participants with hippocampal units completing the memory-based DM task, hippocampal unit analyses were restricted to the value-based DM task. Clustering of hippocampal units otherwise followed the same procedure as for PCC LFPs and units.

### Across task single-unit response comparison

PCC units from the value-based task were clustered into three response profiles, with two characterized by increased peak firing rates (positive units) and one by a decreased peak firing rate (negative units). Units from the memory-based task yielded 4 response profiles, with one characterized by increased peak firing rates, one by a decreased peak, one that did not demonstrate significant peaks, and one excluded due to insufficient membership. To assess whether units demonstrated similar valence (positive/negative) of responses across tasks, we compared whether cluster membership type was matched (e.g. value-task positive cluster and memory-task positive cluster), inverted (e.g. value-task positive cluster and memory-task negative cluster), or other (e.g. value-task positive cluster and memory-task other), and the proportion of units in each category was calculated (Figure 3e-top). To quantify cross-task response magnitude relationships, baseline-corrected firing rates for units identified in both tasks were averaged across trials to yield mean by-block time series separately for trial onset, response, and feedback periods. Each by-unit time series was then smoothed using a 50ms window Gaussian kernel (via ksmooth in R). Linear mixed-effect regression predicting memory-based from value-based responses was carried out separately for congruent, incongruent, and other unit types, including a random intercept term for participant. Pseudo-R^2^ values (marginal coefficient of determination) were calculated using the Multi-Model Inference package (MuMIn, Nakagawa et al., 2017) in R.

### Statistical analyses

BBG and single-unit activity were analyzed using linear mixed-effect models in R (Posit team; R Core Team, 2021) with the lmer (Bates et al., 2015) and lmerTest (Kuznetsova et al., 2017) packages. For group level analyses, each recording site contributed one value, with participant modeled as a random intercept to account for nesting of multiple sites within individuals. Confidence intervals were estimated using Wald estimation. For value- and memory-based DM task trial period analysis, mean activity was computed over 500 ms periods. Primarily these periods were set at: 200 – 700 ms following trial onset and 0 – 500 ms following decision button-press responses and feedback presentation events. For the incidental encoding task, activity was analyzed for two 500 ms periods: 200 – 700 ms following trial onset and −250 – 250 ms centered on 1-back target responses. For the delayed recognition task, activity was analyzed over two 500 ms periods: 200 – 700 ms following trial onset and 0 – 500 ms following the confidence prompt onset. Baseline was considered to be 0 for statistical comparisons. For across task LFP response comparisons, linear mixed-effect regression predicting value-based from memory-based responses, including a random intercept term for participant, was carried out separately for type 1 and type 2 recording sites at each trial time period.

Subjective value of the risky option (SV_risky_) was estimated using an expected utility model, a power function of reward value and reward probability (Jung et al., 2018). Using choice data pooled across all blocks, we estimated two parameters per participant, a risk tolerance parameter (α) and a scaling parameter (β), by fitting a logistic regression that assumes choice depends on the difference in subjective value between the risky and safe options. To quantify the relationship between neural responses and subjective value, mean BBG activity (percent signal change) was extracted from 500 ms onset, response, and feedback time windows as above, and regressed against SV_risky_ values using a linear mixed effect model (BBG ∼ SV_risky_ + (1 | participant/electrode) + (1 | participant:block)), controlling for multiple electrodes within participants as well as for multiple blocks per participant.

In order to evaluate extended neural responses outside of a priori-determined time windows and to control for multiple comparisons across time, we also computed a cluster-mass FWER corrected p-value for all analyses. For each analysis, we divided the timeseries into 500ms windows with 50ms steps, and performed the analysis within each window. Cluster mass served as the test statistic, integrating effect magnitude and duration across windows. Individual windows exceeding a threshold of p < 0.05 were grouped into contiguous clusters, and cluster mass was defined as the sum of absolute t-scores across all windows within a contiguous cluster. A null distribution was constructed by generating 10,000 surrogate timeseries with the same length and autocorrelation as the observed timeseries, and the maximum cluster mass was calculated for each surrogate. Cluster-level, FWER corrected p-values were computed as the proportion of maxima greater than or equal to the observed cluster mass. Where significant clusters overlapped with a priori windows, the corresponding effect is considered supported by the FWER correction. Where an effect identified in a priori windows did not survive cluster correction, corrected p-values are reported in the text.

## Acknowledgments

The authors thank the patients who participated in these experiments and the staff at the Hospital of the University of Pennsylvania, Medical College of Wisconsin, and Baylor St. Luke’s Hospital Epilepsy Monitoring Units for their assistance.

## Author contributions

Conceptualization: B.F., S.K., B.H., J.K.; Experimental procedures: H.C., K.D., S.S., N.P., S. H.; Data collection: S.K.; Data analysis: S.K., B.F., H.R.; Writing – Original Draft: S.K., B.F.; Writing – Review & Editing: all.

