## Supplement for "Electrophysiological dissociation of human posterior cingulate cortex contributions to value- and memory-based decision-making"

Koslov et al.

### Supplementary Results:

| Participant Demographics | | | | | | Electrode Coverage | | | | Behavioral Tasks Completed (# of Blocks) | | | |
|---|---|---|---|---|---|---|---|---|---|---|---|---|---|
| **ID** | **Site** | **Age (yr)** | **Sex** | **Hand** | **Hemisphere** | **PCC Macro** | **PCC Micro** | **Hipp Macro** | **Hipp Micro** | **Value DM** | **Memory DM** | **Encoding** | **Recognition** |
| **PCC Cohort (n = 21)** | | | | | | | | | | | | | |
| **P1** | BCM | 25 | F | R | Bilateral | ✓ | ✓ | ✓ | ✓ | ✓ (4) | — | — | — |
| **P2** | BCM | 38 | M | L | Bilateral | ✓ | ✓ | ✓ | ✓ | ✓ (4) | — | — | — |
| **P3** | MCW | 44 | F | R | Bilateral | ✓ | ✓ | — | ✓ | ✓ (4) | ✓ (2) | ✓ (1) | — |
| **P4** | MCW | 21 | F | L | R | ✓ | ✓ | — | — | ✓ (4) | ✓ (2) | ✓ (1) | ✓ (2) |
| **P5** | Penn | 34 | F | R | R | ✓ | — | — | — | ✓ (4) | — | — | — |
| **P6** | Penn | 59 | F | R | L | ✓ | — | ✓ | — | ✓ (4) | ✓ (2) | ✓ (1) | ✓ (2) |
| **P7** | Penn | 28 | F | R | Bilateral | ✓ | — | ✓ | — | ✓ (2) | ✓ (2) | ✓ (1) | ✓ (2) |
| **P8** | Penn | 29 | F | R | Bilateral | ✓ | — | ✓ | — | ✓ (2) | ✓ (2) | ✓ (1) | — |
| **P9** | Penn | 34 | M | R | L | ✓ | — | — | — | ✓ (2) | ✓ (2) | ✓ (1) | — |
| **P10** | Penn | 26 | M | L | L | ✓ | — | ✓ | — | ✓ (2) | — | — | — |
| **P11** | Penn | 61 | F | R | Bilateral | ✓ | ✓ | ✓ | — | ✓ (3) | ✓ (2) | ✓ (1) | ✓ (2) |
| **P12** | Penn | 34 | F | R | R | ✓ | — | ✓ | — | ✓ (4) | ✓ (2) | ✓ (1) | ✓ (2) |
| **P13** | Penn | 46 | F | R | L | ✓ | — | ✓ | ✓ | ✓ (4) | ✓ (2) | ✓ (1) | ✓ (2) |
| **P14** | Penn | 57 | F | R | L | ✓ | ✓ | — | — | ✓ (4) | ✓ (2) | ✓ (1) | ✓ (2) |
| **P15** | Penn | 33 | M | R | Bilateral | ✓ | — | ✓ | — | ✓ (4) | ✓ (2) | ✓ (1) | ✓ (2) |
| **P16** | Penn | 47 | M | R | Bilateral | ✓ | ✓ | — | — | ✓ (4) | ✓ (2) | ✓ (1) | ✓ (2) |
| **P17** | Penn | 32 | M | R | L | ✓ | — | — | — | ✓ (2) | ✓ (2) | ✓ (1) | — |
| **P18** | Penn | 29 | M | R | L | ✓ | ✓ | ✓ | ✓ | ✓ (4) | ✓ (2) | ✓ (1) | ✓ (2) |
| **P19** | Penn | 47 | M | R | Bilateral | ✓ | — | ✓ | ✓ | ✓ (4) | ✓ (2) | — | ✓ (2) |
| **P20** | Penn | 20 | M | R | L | ✓ | ✓ | — | ✓ | ✓ (4) | — | ✓ (1) | ✓ (2) |
| **P38** | MCW | 55 | F | R | Bilateral | — | ✓ | — | ✓ | ✓ (4) | — | — | — |
| **Hippocampal Cohort (n = 17)** | | | | | | | | | | | | | |
| **P21** | BCM | 68 | M | R | Bilateral | — | — | ✓ | ✓ | ✓ (4) | — | — | — |
| **P22** | BCM | 22 | M | R | Bilateral | — | — | ✓ | ✓ | ✓ (4) | — | — | — |
| **P23** | BCM | 28 | M | L | Bilateral | — | — | ✓ | ✓ | ✓ (2) | — | — | — |
| **P24** | BCM | 27 | F | L | Bilateral | — | — | — | ✓ | ✓ (2) | — | — | — |
| **P25** | BCM | 37 | F | R | Bilateral | — | — | — | ✓ | ✓ (2) | — | — | — |
| **P26** | BCM | 20 | F | R | Bilateral | — | — | ✓ | ✓ | ✓ (4) | ✓ (2) | ✓ (1) | ✓ (1) |
| **P27** | BCM | 41 | F | R | Bilateral | — | — | — | ✓ | ✓ (4) | — | — | — |
| **P28** | Penn | 31 | M | R | Bilateral | — | — | ✓ | — | ✓ (3) | ✓ (2) | ✓ (1) | ✓ (2) |
| **P29** | Penn | 57 | M | R | Bilateral | — | — | ✓ | — | ✓ (2) | — | — | — |
| **P30** | Penn | 26 | M | L | L | — | — | ✓ | — | ✓ (3) | ✓ (2) | ✓ (1) | ✓ (2) |
| **P31** | Penn | 28 | F | R | Bilateral | — | — | ✓ | — | ✓ (4) | ✓ (2) | ✓ (1) | ✓ (1) |
| **P32** | Penn | 46 | M | R | Bilateral | — | — | ✓ | ✓ | ✓ (3) | ✓ (2) | ✓ (1) | — |
| **P33** | Penn | 64 | M | R | R | — | — | ✓ | — | ✓ (2) | — | — | — |
| **P34** | Penn | 21 | F | R | Bilateral | — | — | ✓ | ✓ | ✓ (2) | ✓ (2) | ✓ (1) | — |
| **P35** | Penn | 44 | F | L | L | — | — | ✓ | ✓ | ✓ (4) | ✓ (2) | ✓ (1) | ✓ (2) |
| **P36** | Penn | 29 | M | R | Bilateral | — | — | ✓ | ✓ | ✓ (4) | ✓ (2) | ✓ (1) | ✓ (2) |
| **P37** | Penn | 31 | M | R | Bilateral | — | — | ✓ | — | ✓ (2) | ✓ (2) | ✓ (1) | ✓ (2) |

**Table S1. Participant information**. Details of participant macroelectrode and microwire recording locations, clinical recording site, task completion, and demographics. Check marks indicate that a participant had macroelectrode or microwire recordings or completed a given task; numbers in parentheses indicate the number of task blocks completed, and a dash (—) indicates no recording or data. *Abbreviations:* Penn: Hospital of the University of Pennsylvania; BCM: Baylor St. Luke's Hospital / Baylor College of Medicine; MCW: Medical College of Wisconsin; PCC: posterior cingulate cortex; Hipp: hippocampus; Macro: clinical macroelectrode; Micro: microwire array; F: female; M: male; Hand: handedness (R, right; L, left); Hemi: hemisphere of implant (R, right; L, left; Bilateral, both hemispheres); DM: decision-making; Encoding: incidental encoding 1-back task; Recognition: delayed recognition memory task.

| PCC Decision-Making Macro Electrode Anatomical Statistics Reporting Table | | | | | | |
|---|---|---|---|---|---|---|
| **Category** | **Comparison** | **Task** | **Time** | **β-value [95% CI]** | **t-score (df)** | **p-value** |
| dPCC | vs. Baseline | Value-based | Onset | 6.371 [3.415, 9.328] | 4.224 (14.8) | < 0.001 *** |
| | | | Response | 5.245 [2.922, 7.568] | 4.426 (14.3) | < 0.001 *** |
| | | | Feedback | 2.694 [1.755, 3.633] | 5.623 (13.2) | < 0.001 *** |
| | | Memory-based | Onset | 9.012 [4.269, 13.756] | 3.724 (10.1) | 0.004 ** |
| | | | Response | 3.906 [1.534, 6.273] | 3.234 (10.2) | 0.008 ** |
| | | | Feedback | 2.938 [1.361, 4.516] | 3.652 (10.1) | 0.004 ** |
| vPCC | vs. Baseline | Value-based | Onset | 1.949 [0.323, 3.575] | 2.349 (11.9) | 0.037 * |
| | | | Response | 0.610 [-0.928, 2.148] | 0.777 (12.9) | 0.451 |
| | | | Feedback | 0.817 [0.022, 1.612] | 2.014 (13.6) | 0.064 |
| | | Memory-based | Onset | 1.756 [-0.513, 4.025] | 1.517 (8.0) | 0.168 |
| | | | Response | 0.860 [-1.359, 3.080] | 0.760 (6.8) | 0.473 |
| | | | Feedback | 1.313 [-0.279, 2.906] | 1.616 (8.0) | 0.145 |

**Table S2. PCC decision-making task statistics for anatomically labeled electrode groups**. Statistics are shown for linear mixed-effects regression for dPCC and vPCC recording macroelectrode sites against baseline for the value-based and memory-based tasks across onset, response, and feedback time periods. β-value = regression coefficient, df = degrees of freedom estimated using Satterthwaite's approximation. *$p < 0.05$, **$p < 0.01$, ***$p < 0.001$.

| PCC Decision-Making Task Macro Electrode Cluster Statistics Reporting Table | | | | | | |
|---|---|---|---|---|---|---|
| **Table S3a - Main Effect** | | | | | | |
| **Category** | **Comparison** | **Task** | **Time** | **β-value [95% CI]** | **t-score (df)** | **p-value** |
| Cluster Type 1 | vs. Baseline | Value-based | Onset | 6.893 [4.642, 9.143] | 6.003 (18.2) | < 0.001 *** |
| | | | Response | 5.385 [3.613, 7.156] | 5.958 (17.6) | < 0.001 *** |
| | | | Feedback | 2.790 [1.944, 3.635] | 6.468 (14.1) | < 0.001 *** |
| | | Memory-based | Onset | 9.039 [5.019, 13.060] | 4.407 (9.5) | 0.001 ** |
| | | | Response | 4.780 [3.228, 6.331] | 6.038 (9.3) | < 0.001 *** |
| | | | Feedback | 3.400 [2.291, 4.509] | 6.008 (8.0) | < 0.001 *** |
| Cluster Type 2 | vs. Baseline | Value-based | Onset | -0.165 [-1.512, 1.183] | -0.239 (6.2) | 0.819 |
| | | | Response | -1.145 [-2.416, 0.126] | -1.765 (6.5) | 0.124 |
| | | | Feedback | 0.301 [-0.686, 1.287] | 0.597 (7.0) | 0.569 |
| | | Memory-based | Onset | 0.735 [-2.186, 3.656] | 0.493 (5.2) | 0.642 |
| | | | Response | -1.406 [-3.255, 0.443] | -1.490 (4) | 0.210 |
| | | | Feedback | -0.271 [-1.773, 1.231] | -0.354 (5.4) | 0.737 |
| **Table S3b - Risk Modulation** | | | | | | |
| Cluster Type 1 | Risky > Safe | Value-based | Onset | 0.032 [-1.149, 1.215] | 0.054 (15.9) | 0.958 |
| | | | Response | 3.149 [1.427, 4.871] | 3.584 (16.1) | 0.002 ** |
| | | | Feedback | 0.966 [-0.484, 2.415] | 1.306 (15.4) | 0.211 |
| | | Memory-based | Onset | 0.915 [-1.149, 3.315] | 0.747 (8.1) | 0.476 |
| | | | Response | 2.941 [0.560, 5.321] | 2.421 (9.4) | 0.038 * |
| | | | Feedback | 2.390 [0.329, 4.450] | 2.273 (10.2) | 0.046 * |
| Cluster Type 2 | Risky > Safe | Value-based | Onset | 1.757 [-1.400, 4.915] | 1.091 (4.5) | 0.331 |
| | | | Response | 0.210 [-2.237, 2.692] | 0.166 (5.8) | 0.874 |
| | | | Feedback | 0.334 [-2.938, 3.607] | 0.200 (6.1) | 0.848 |
| | | Memory-based | Onset | 1.594 [-0.139, 3.327] | 1.803 (14) | 0.093 |
| | | | Response | 1.673 [-0.122, 3.468] | 1.826 (4.3) | 0.137 |
| | | | Feedback | 1.197 [-0.558, 2.952] | 1.337 (4.4) | 0.246 |
| **Table S3c - Subjective Value** | | | | | | |
| Cluster Type 1 | Subjective Value | Value-based | Onset | 0.041 [-0.018, 0.100] | 1.373 (2194.0) | 0.17 |
| | | | Response | 0.091 [0.034, 0.149] | 3.113 (1180.7) | 0.002 ** |
| | | | Feedback | 0.052 [0.001, 0.103] | 1.940 (189.9) | 0.053 |
| Cluster Type 2 | | | Onset | -0.009 [-0.104, 0.086] | -0.183 (85.0) | 0.855 |
| | | | Response | -0.014 [-0.110, 0.082] | -0.284 (390.6) | 0.776 |
| | | | Feedback | 0.024 [-0.067, 0.115] | 0.517 (46.5) | 0.608 |
| **Table S3d - Across Task Comparison** | | | | | | |
| Cluster Type 1 | Across-task correlation | | Onset | 0.625 [0.439, 0.811] | 6.589 (15) | < 0.001 *** |
| | | | Response | 1.441 [0.973, 1.907] | 6.046 (9.6) | < 0.001 *** |
| | | | Feedback | 0.411 [-0.022, 0.844] | 1.861 (15) | 0.083 |
| Cluster Type 2 | Across-task correlation | | Onset | 0.062 [-0.145, 0.268] | 0.586 (9) | 0.572 |
| | | | Response | 0.345 [-0.114, 0.805] | 1.473 (12.5) | 0.165 |
| | | | Feedback | 0.219 [-0.110, 0.548] | 1.304 (15) | 0.216 |

**Table S3. PCC decision-making statistics for clustered macroelectrode recording sites**. Table S3a shows the results from linear mixed-effects regression comparing BBG percent signal change against baseline for cluster type 1 and cluster type 2 macroelectrode recording sites from the value-based and memory-based decision-making tasks during the onset, response, and feedback time periods. Table S3b shows regression results comparing responses from trials where risky versus safe decisions were made for each time period. Table S3c shows regression result comparing cluster responses against subjective value estimates across value-based task time periods. Table S3d shows the comparison across the value- and memory-based decision-making tasks of average percent signal change responses from cluster types during each time period. See methods for more details. β-value = regression coefficient, df = degrees of freedom estimated using Satterthwaite's approximation. *$p < 0.05$, **$p < 0.01$, ***$p < 0.001$.

| PCC Single-Unit Decision-Making Statistics Reporting Table | | | | | | |
|---|---|---|---|---|---|---|
| **Table S4a - Main Effect** | | | | | | |
| **Category** | **Comparison** | **Task** | **Time** | **β-value [95% CI]** | **t-score (df)** | **p-value** |
| Unit Type 1 | vs. Baseline | Value-based | Onset | -0.454 [-1.039, 0.130] | -1.523 (27) | 0.139 |
| | | | Response | 0.587 [-0.199, 1.372] | 1.464 (5.7) | 0.196 |
| | | | Feedback | 0.776 [0.060, 1.491] | 2.123 (5.3) | 0.084 |
| | | | Feedback-Centered | 1.186 [0.360, 2.011] | 2.816 (5.4) | 0.034 * |
| Unit Type 2 | | | Onset | 1.092 [0.237, 1.947] | 2.504 (9.4) | 0.032 * |
| | | | Response | 0.404 [-0.245, 1.052] | 1.220 (7.3) | 0.26 |
| | | | Feedback | -0.130 [-0.578, 0.318] | -0.570 (6.9) | 0.587 |
| Unit Type 3 | | | Onset | -0.570 [-0.998, -0.142] | -2.608 (6.4) | 0.038 * |
| | | | Response | -1.018 [-1.536, -0.500] | -3.852 (8.5) | 0.004 ** |
| | | | Feedback | -0.299 [-0.786, 0.187] | -1.205 (6.4) | 0.271 |
| Unit Type 1 | | Memory-based | Onset | 2.680 [1.066, 4.294] | 3.255 (3.7) | 0.035 * |
| | | | Response | 0.905 [0.119, 1.691] | 2.257 (3.3) | 0.101 |
| | | | Feedback | 0.458 [-0.029, 0.946] | 1.844 (4.7) | 0.129 |
| Unit Type 2 | | | Onset | -1.349 [-2.170, -0.527] | -3.217 (4.3) | 0.029 * |
| | | | Response | -0.827 [-2.119, 0.466] | -1.254 (4.4) | 0.272 |
| | | | Feedback | 0.059 [-1.359, 1.478] | 0.082 (4.5) | 0.938 |
| Unit Type 3 | | | Onset | -0.221 [-0.701, 0.259] | -0.903 (2.1) | 0.457 |
| | | | Response | -0.175 [-0.709, 0.359] | -0.643 (1.3) | 0.615 |
| | | | Feedback | 0.105 [-0.231, 0.440] | 0.611 (4) | 0.574 |

| Table S4b - Risk Modulation | | | | | | |
|---|---|---|---|---|---|---|
| Unit Type 1 | Risky > Safe | Value-based | Onset | -0.128 [-0.691, 0.435] | -0.445 (2.6) | 0.691 |
| | | | Response | 0.660 [-0.193, 1.513] | 1.517 (27) | 0.141 |
| | | | Feedback | 0.762 [0.087, 1.436] | 2.213 (27) | 0.035 * |
| | | | Feedback-Centered | 1.068 [0.225, 1.909] | 2.483 (27) | 0.02 * |
| Unit Type 2 | | | Onset | 0.121 [-0.423, 0.664] | 0.434 (35) | 0.667 |
| | | | Response | 0.062 [-0.955, 1.078] | 0.119 (8.6) | 0.908 |
| | | | Feedback | 0.001 [-0.724, 0.726] | 0.003 (5.6) | 0.998 |
| Unit Type 3 | | | Onset | -0.313 [-0.768, 0.141] | -1.351 (3.7) | 0.254 |
| | | | Response | -0.420 [-0.986, 0.146] | -1.454 (4.2) | 0.216 |
| | | | Feedback | 0.140 [-0.251, 0.530] | 0.701 (45) | 0.487 |
| Unit Type 1 | | Memory-based | Onset | 0.229 [-0.385, 0.843] | 0.731 (0.7) | 0.634 |
| | | | Response | -0.200 [-1.095, 0.696] | -0.437 (5.2) | 0.68 |
| | | | Feedback | 0.281 [-1.085, 1.646] | 0.403 (5.0) | 0.704 |
| Unit Type 2 | | | Onset | 0.039 [-0.489, 0.567] | 0.146 (3.7) | 0.891 |
| | | | Response | 0.084 [-0.679, 0.848] | 0.217 (3.5) | 0.84 |
| | | | Feedback | 0.800 [0.153, 1.446] | 2.425 (28) | 0.022 * |
| Unit Type 3 | | | Onset | 2.074 [-1.309, 5.457] | 1.202 (1.8) | 0.362 |
| | | | Response | 1.278 [-0.683, 3.238] | 1.277 (2.1) | 0.325 |
| | | | Feedback | 1.342 [-1.351, 4.036] | 0.977 (2.2) | 0.424 |

| Table S4c - PCC Across Decision-Making Tasks Unit Type Comparison | | | | | | |
|---|---|---|---|---|---|---|
| **Category** | **Comparison** | **Time** | **r-squared** | **β-value [95% CI]** | **t-score (df)** | **p-value** |
| Congruent | Across-Task | Onset | 0.018 | 0.181 [-0.357, 0.720] | 0.660 (16.2) | 0.519 |
| | | Response | 0.439 | 0.849 [0.455, 1.243] | 4.226 (16.9) | < 0.001 *** |
| | | Feedback | 0.187 | 0.458 [0.092, 0.824] | 2.451 (16.2) | 0.022 * |
| Incongruent | | Onset | 0.295 | -1.074 [-1.835, -0.312] | -2.764 (13.4) | 0.016 * |
| | | Response | 0.142 | 0.379 [-0.097, 0.854] | 1.562 (6.4) | 0.166 |
| | | Feedback | 0.274 | 0.926 [0.326, 1.527] | 3.024 (13.3) | 0.010 * |
| Other | | Onset | 0.063 | 0.235 [-0.490, 0.960] | 0.634 (5) | 0.554 |
| | | Response | 0.092 | 0.237 [-0.360, 0.834] | 0.779 (5) | 0.471 |
| | | Feedback | 0.108 | 0.307 [-0.400, 1.015] | 0.851 (5) | 0.433 |

**Table S4. PCC microwire decision-making statistics for unit-type clusters**. Table S4a shows results from linear mixed-effects regression comparing normalized firing rates to baseline for PCC unit types from the value-based and memory-based decision-making tasks during all time periods. Table S4b shows results of regressions comparing adjusted firing rate from trials with risky versus safe decisions for clusters across time periods from each task. Table S4c shows the comparison of average firing rate across the value-based and memory-based decision-making tasks during each time period for units identified as congruent, incongruent, or other. See methods for more details. β-value = regression coefficient, df = degrees of freedom estimated using Satterthwaite's approximation. *p < 0.05, **p < 0.01, ***p < 0.001.

| Hippocampus Decision-Making Macro Electrode Anatomical Statistics Reporting Table | | | | | | |
|---|---|---|---|---|---|---|
| **Table S5a - Hippocampus Main Effect** | | | | | | |
| **Category** | **Comparison** | **Task** | **Time** | **β-value [95% CI]** | **t-score (df)** | **p-value** |
| Anterior Hipp. | vs. Baseline | Value-based | Onset | 1.731 [1.210, 2.252] | 6.511 (23.7) | < 0.001 *** |
| | | | Response | 0.579 [-0.206, 1.363] | 1.446 (19.1) | 0.164 |
| | | | Feedback | 1.023 [0.440, 1.605] | 3.439 (20.3) | 0.003 ** |
| Posterior Hipp. | | | Onset | 2.361 [1.498, 3.224] | 5.364 (19.5) | < 0.001 *** |
| | | | Response | 1.333 [0.309, 2.357] | 2.551 (20.3) | 0.019 * |
| | | | Feedback | 2.064 [1.287, 2.841] | 5.205 (21.1) | < 0.001 *** |
| Anterior Hipp. | | Memory-based | Onset | 4.106 [2.913, 5.300] | 6.743 (13.8) | < 0.001 *** |
| | | | Response | 1.177 [0.446, 1.908] | 3.156 (13.8) | 0.006 ** |
| | | | Feedback | 1.185 [0.312, 2.060] | 2.660 (14.7) | 0.018 * |
| Posterior Hipp. | | | Onset | 2.970 [1.716, 4.223] | 4.644 (14.2) | < 0.001 *** |
| | | | Response | 1.445 [0.307, 2.583] | 2.488 (13.6) | 0.027 * |
| | | | Feedback | 1.704 [0.731, 2.680] | 3.434 (12.5) | 0.005 ** |
| **Table S5b - Anterior Vs. Posterior, BBG Response** | | | | | | |
| Ant. vs. Post. Hipp. | Main Effect | Value-based | Onset | 0.770 [0.203, 1.338] | 2.661 (129.3) | 0.009 ** |
| | | | Response | 1.065 [0.369, 1.762] | 2.997 (129.3) | 0.003 ** |
| | | | Feedback | 1.428 [0.929, 1.927] | 5.610 (124.6) | < 0.001 *** |
| | | Memory-based | Onset | -0.606 [-1.597, 0.385] | -1.199 (84.6) | 0.234 |
| | | | Response | 0.335 [-0.570, 1.238] | 0.726 (89.8) | 0.47 |
| | | | Feedback | 0.683 [-0.144, 1.509] | 1.618 (88.0) | 0.109 |
| **Table S5c - Hippocampus Risk Modulation** | | | | | | |
| Anterior Hipp. | Risky > Safe | Value-based | Onset | 0.284 [-0.611, 1.180] | 0.622 (22.4) | 0.540 |
| | | | Response | -0.095 [-0.927, 0.737] | -0.224 (22.3) | 0.825 |
| | | | Feedback | -0.640 [-1.635, 0.355] | -1.260 (23.2) | 0.220 |
| Posterior Hipp. | | | Onset | 0.975 [-0.007, 1.957] | 1.946 (20.7) | 0.065 |
| | | | Response | 1.885 [0.743, 3.028] | 3.234 (19.6) | 0.004 ** |
| | | | Feedback | 0.729 [-0.111, 1.569] | 1.702 (14.5) | 0.110 |
| Anterior Hipp. | | Memory-based | Onset | 0.948 [-0.138, 2.035] | 1.711 (13.9) | 0.109 |
| | | | Response | 0.814 [-0.211, 1.839] | 1.556 (56) | 0.125 |
| | | | Feedback | -0.816 [-2.165, 0.534] | -1.184 (14.2) | 0.256 |
| Posterior Hipp. | | | Onset | 0.190 [-1.108, 1.489] | 0.287 (12.9) | 0.779 |
| | | | Response | 2.232 [0.054, 4.409] | 2.009 (14.9) | 0.063 |
| | | | Feedback | 1.669 [0.339, 2.998] | 2.460 (13.0) | 0.029 * |

**Table S5. Hippocampal macroelectrode statistics table for decision-making tasks**. Table S5a shows results from linear mixed-effects regression comparing BBG percent signal change against baseline from the value-based and memory-based decision-making tasks across all time periods. Table S5b shows the comparison of response magnitude between anterior versus posterior recording sites during the decision-making tasks from all time periods. Table S5c shows results of regression comparing responses from trials with risky versus safe decisions across tasks and time periods. β-value = regression coefficient, df = degrees of freedom estimated using Satterthwaite's approximation. *p < 0.05, **p < 0.01, ***p < 0.001.

| 1-back Incidental Encoding All Regions Statistics Reporting Table | | | | | | |
|---|---|---|---|---|---|---|
| **Table S6a - Trial Onset** | | | | | | |
| **Category** | **Comparison** | **Task** | **Time** | **β-value [95% CI]** | **t-score (df)** | **p-value** |
| dPCC | vs. Baseline | Encoding | Trial Onset | 5.778 [3.070, 8.486] | 4.182 (8.6) | 0.003 ** |
| vPCC | | | | 1.210 [-0.367, 2.787] | 1.504 (7.0) | 0.176 |
| Anterior Hipp. | | | | 3.842 [3.032, 4.651] | 9.302 (13.9) | < 0.001 *** |
| Posterior Hipp. | | | | 2.989 [1.955, 4.023] | 5.665 (11.5) | < 0.001 *** |
| vPCC | vs. dPCC | Encoding | Trial Onset | -4.996 [-6.664, -3.327] | -5.870 (100.5) | < 0.001 *** |
| Anterior Hipp. | | | | -2.284 [-3.797, -0.770] | -2.957 (105.9) | 0.003 ** |
| Posterior Hipp. | | | | -2.988 [-4.551, -1.424] | -3.746 (106.9) | < 0.001 *** |
| **Table S6b - Target Onset** | | | | | | |
| dPCC | vs. Baseline | Encoding | Target Onset | 12.170 [5.474, 18.866] | 3.562 (10.0) | 0.005 ** |
| vPCC | | | | -0.728 [-3.634, 2.178] | -0.491 (6.5) | 0.639 |
| Anterior Hipp. | | | | 4.537 [1.774, 7.301] | 3.218 (11.5) | 0.008 ** |
| Posterior Hipp. | | | | 6.447 [3.980, 8.911] | 5.127 (11.9) | < 0.001 *** |
| vPCC | vs. dPCC | Encoding | Target Onset | -11.382 [-14.627, -8.137] | -6.875 (96.2) | < 0.001 *** |
| Anterior Hipp. | | | | -7.096 [-10.208, -3.984] | -4.469 (105.2) | < 0.001 *** |
| Posterior Hipp. | | | | -4.597 [-7.781, -1.413] | -2.830 (104.2) | 0.006 ** |

**Table S6. 1-back incidental encoding task statistics**. Table S6a-top shows the results of linear mixed-effects regression comparing BBG percent signal change responses to baseline from all subregions (dPCC, vPCC, anterior hippocampus, and posterior hippocampus) during the trial onset period of the incidental encoding task. Table S6a-bottom shows the results of regressions comparing dPCC to all other subregions during the trial onset period of the incidental encoding task. Table S6b-top shows the results of linear mixed-effects regression comparing percent signal change responses to baseline from all subregions for the time period centered around 1-back target responses during the incidental encoding task. Table S6b-bottom shows the results of regressions comparing dPCC to all other subregions during the target period. β-value = regression coefficient, df = degrees of freedom estimated using Satterthwaite's approximation. *$p < 0.05$, **$p < 0.01$, ***$p < 0.001$.

| Delayed Recognition All Regions Statistics Reporting Table | | | | | | |
|---|---|---|---|---|---|---|
| **Table S7a - Trial Onset** | | | | | | |
| **Category** | **Comparison** | **Task** | **Time** | **β-value [95% CI]** | **t-score (df)** | **p-value** |
| dPCC | vs. Baseline | Delayed Rec. | Trial Onset | 6.099 [2.614, 9.584] | 3.430 (13) | 0.004 ** |
| vPCC | | | | 2.279 [-1.423, 5.981] | 1.206 (6.2) | 0.272 |
| ant hipp | | | | 5.970 [4.074, 7.866] | 6.170 (11.5) | < 0.001 *** |
| post hipp | | | | 2.689 [0.972, 4.404] | 3.071 (10.2) | 0.012 * |
| vPCC | vs. dPCC | Delayed Rec. | Trial Onset | -5.084 [-8.057, -2.112] | -3.352 (97.0) | 0.001 ** |
| ant hipp | | | | -1.410 [-4.111, 1.291] | -1.023 (99.2) | 0.309 |
| post hipp | | | | -4.213 [-7.022, -1.404] | -2.940 (99.9) | 0.004 ** |
| **Table S7b - Confidence Judgments** | | | | | | |
| dPCC | vs. Baseline | Delayed Rec. | Confidence Onset | 2.642 [0.490, 4.793] | 2.407 (13) | 0.032 * |
| vPCC | | | | -0.662 [-1.837, 0.513] | -1.104 (6.0) | 0.311 |
| ant hipp | | | | -0.231 [-0.959, 0.497] | -0.622 (10.7) | 0.547 |
| post. hipp | | | | -0.950 [-1.864, -0.036] | -2.038 (11.1) | 0.066 |
| vPCC | vs. dPCC | Delayed Rec. | Confidence Onset | -3.490 [-4.918, -2.062] | -4.789 (97.5) | < 0.001 *** |
| ant. Hipp | | | | -3.205 [-4.496, -1.914] | -4.865 (98.3) | < 0.001 *** |
| post. Hipp | | | | -3.666 [-5.011, -2.322] | -5.345 (99.6) | < 0.001 *** |

**Table S7. Delayed recognition task statistics**. Table S7a-top shows the results of linear mixed-effects regression comparing BBG percent signal change responses to baseline from all subregions (dPCC, vPCC, anterior hippocampus, and posterior hippocampus) during the trial onset period for item-recognition decisions of the delayed recognition task. Table S7a-bottom shows the results of regressions comparing dPCC to all other subregions during the trial onset period of the delayed recognition task. Table S7b-top shows the results of linear mixed-effects regression comparing percent signal change responses to baseline from all subregions for the confidence judgment time period of the delayed recognition task. Table S7b-bottom shows the results of regressions comparing dPCC to all other subregions during the confidence judgment period. β-value = regression coefficient, df = degrees of freedom estimated using Satterthwaite's approximation. *p < 0.05, **p < 0.01, ***p < 0.001.

| Hippocampus Single-Unit Decision-Making Statistics Reporting Table | | | | | | |
|---|---|---|---|---|---|---|
| Table S8a - Trial Onset | | | | | | |
| **Category** | **Comparison** | **Task** | **Time** | **β-value [95% CI]** | **t-score (df)** | **p-value** |
| Unit Type 1 | vs. Baseline | Value-based | Onset | 0.512 [-0.242, 1.385] | 1.377 (2.1) | 0.297 |
| | | | Response | -0.365 [-0.795, 0.064] | -1.667 (2.1) | 0.234 |
| | | | Feedback | -1.008 [-2.452, 0.436] | -1.368 (2.1) | 0.299 |
| Unit Type 2 | | | Onset | 0.423 [-0.504, 1.350] | 0.894 (3.936) | 0.422 |
| | | | Response | 0.986 [-0.113, 2.085] | 1.758 (5.1) | 0.137 |
| | | | Feedback | 0.237 [-0.224, 0.698] | 1.006 (4.9) | 0.362 |
| | | | Feedback-centered | 0.981 [0.258, 1.703] | 2.660 (4.9) | 0.046 * |
| Unit Type 3 | | | Onset | -1.118 [-1.720, -0.517] | -3.645 (8) | 0.007 ** |
| | | | Response | -2.342 [-3.782, -0.902] | -3.187 (4.9) | 0.026 * |
| | | | Feedback | -0.233 [-1.789, 1.322] | -0.294 (4.9) | 0.779 |
| Table S8b - Risk Modulation | | | | | | |
| Unit Type 1 | Risky > Safe | Value-based | Onset | -0.496 [-2.849, 1.858] | -0.413 (2.0) | 0.720 |
| | | | Response | -0.576 [-3.771, 2.619] | -0.353 (1.9) | 0.759 |
| | | | Feedback | -0.512 [-4.439, 3.416] | -0.255 (1.8) | 0.825 |
| Unit Type 2 | | | Onset | -0.006 [-0.471, 0.460] | -0.025 (2.5) | 0.982 |
| | | | Response | 0.407 [-0.602, 1.415] | 0.791 (5.2) | 0.463 |
| | | | Feedback | -0.722 [-2.410, 0.964] | -0.839 (4.4) | 0.445 |
| Unit Type 3 | | | Onset | 0.597 [-0.542, 1.736] | 1.027 (6.1) | 0.343 |
| | | | Response | 0.151 [-0.789, 1.091] | 0.314 (5.9) | 0.764 |
| | | | Feedback | 1.946 [-1.873, 5.766] | 0.999 (5.9) | 0.357 |

**Table S8. Hippocampal microwire value-based task statistics**. Table S8a shows the results of linear mixed-effects regression comparing normalized firing rate responses to baseline during the value-based decision-making task for hippocampal unit cluster types from all time periods. Table S8b shows the results of regressions comparing response magnitude from trials with risky versus safe decisions for cluster types across value-based decision-making task time periods. β-value = regression coefficient, df = degrees of freedom estimated using Satterthwaite's approximation. *p < 0.05, **p < 0.01, ***p < 0.001.
